# Dynamic consensus-building between neocortical areas via long-range connections

**DOI:** 10.1101/2024.11.27.625691

**Authors:** Mitra Javadzadeh, Marine Schimel, Sonja B. Hofer, Yashar Ahmadian, Guillaume Hennequin

## Abstract

The neocortex is organized into functionally specialized areas. While the functions and underlying neural circuitry of individual neocortical areas are well studied, it is unclear how these regions operate collectively to form percepts and implement cognitive processes. In particular, it remains unknown how distributed, potentially conflicting computations can be reconciled. Here we show that the reciprocal excitatory connections between cortical areas orchestrate neural dynamics to facilitate the gradual emergence of a ‘consensus’ across areas. We investigated the joint neural dynamics of primary (V1) and higher-order lateromedial (LM) visual areas in mice, using simultaneous multi-area electrophysiological recordings along with focal optogenetic perturbations to causally manipulate neural activity. We combined mechanistic circuit modeling with state-of-the-art data-driven nonlinear system identification, to construct biologically-constrained latent circuit models of the data that we could further interrogate. This approach revealed that long-range, reciprocal excitatory connections between V1 and LM implement an approximate line attractor in their joint dynamics, which promotes activity patterns encoding the presence of the stimulus consistently across the two areas. Further theoretical analyses revealed that the emergence of line attractor dynamics is a signature of a more general principle governing multi-area network dynamics: reciprocal inter-area excitatory connections reshape the dynamical landscape of the network, specifically slowing down the decay of activity patterns that encode stimulus features congruently across areas, while accelerating the decay of inconsistent patterns. This selective dynamic amplification leads to the emergence of multi-dimensional consensus between cortical areas about various stimulus features. Our analytical framework further predicted the timescales of specific activity patterns across areas, which we directly verified in our data. Therefore, by linking the anatomical organization of inter-area connections to the features they reconcile across areas, our work introduces a general theory of multi-area computation.

---

The neocortex is segregated into distinct areas that are specialized for specific functions. This organization allows for decomposing complex problems into simpler sub-computations, such as the extraction of low-level features from intricate visual scenes. However, cognition arises from the holistic integration of these processes, making it essential that the different areas work in concert and remain consistent with each other. It is unclear how such coordination is achieved, and in particular how any conflict that might arise between local subunits can be globally resolved.

Anatomically, cortical areas are densely interconnected through reciprocal long-range inter-area connections [Felleman and Van Essen, 1991], whose organization is markedly distinct from that of local circuits within a cortical area. For instance, both excitatory and inhibitory neurons have local innervation, while only excitatory neurons have long-range projections that may target other areas [Douglas and Martin, 2004, Markram et al., 2004, Harris and Shepherd, 2015]. The functional role of these distinct connectivity rules is not clear; it remains unknown how excitatory inter-area connections coordinate cortical activity and unify local sub-units into coherent global computations. To address this, we combined mechanistic modelling of cortical circuits with data-driven inference of circuit dynamics. This approach allowed us to build models of cortical activity that not only explained neural responses quantitatively, but also captured the causal effects of optogenetic perturbations and had biologically interpretable components – including local and long-range connections – whose functional significance we could interrogate.

We focused on the joint activity dynamics of the primary (V1) and higher-order (LM) visual areas in mice during visual processing. We used simultaneous multi-channel recordings from V1 and LM performed while mice were presented with a 500 ms-long visual stimulus – one of two stationary gratings oriented at 45° or -45°(Figure 1A-B). Mice were trained to perform a go/no-go task, discriminating the two stimuli. In some trials, neural activity in either V1 or LM (varying across animals) was perturbed in brief 150 ms time windows, using optogenetic activation of inhibitory parvalbumin-expressing (PV+) interneurons expressing channelrhodopsin-2 (ChR2) (Figure 1C) [Javadzadeh and Hofer, 2022].

**Figure 1.**
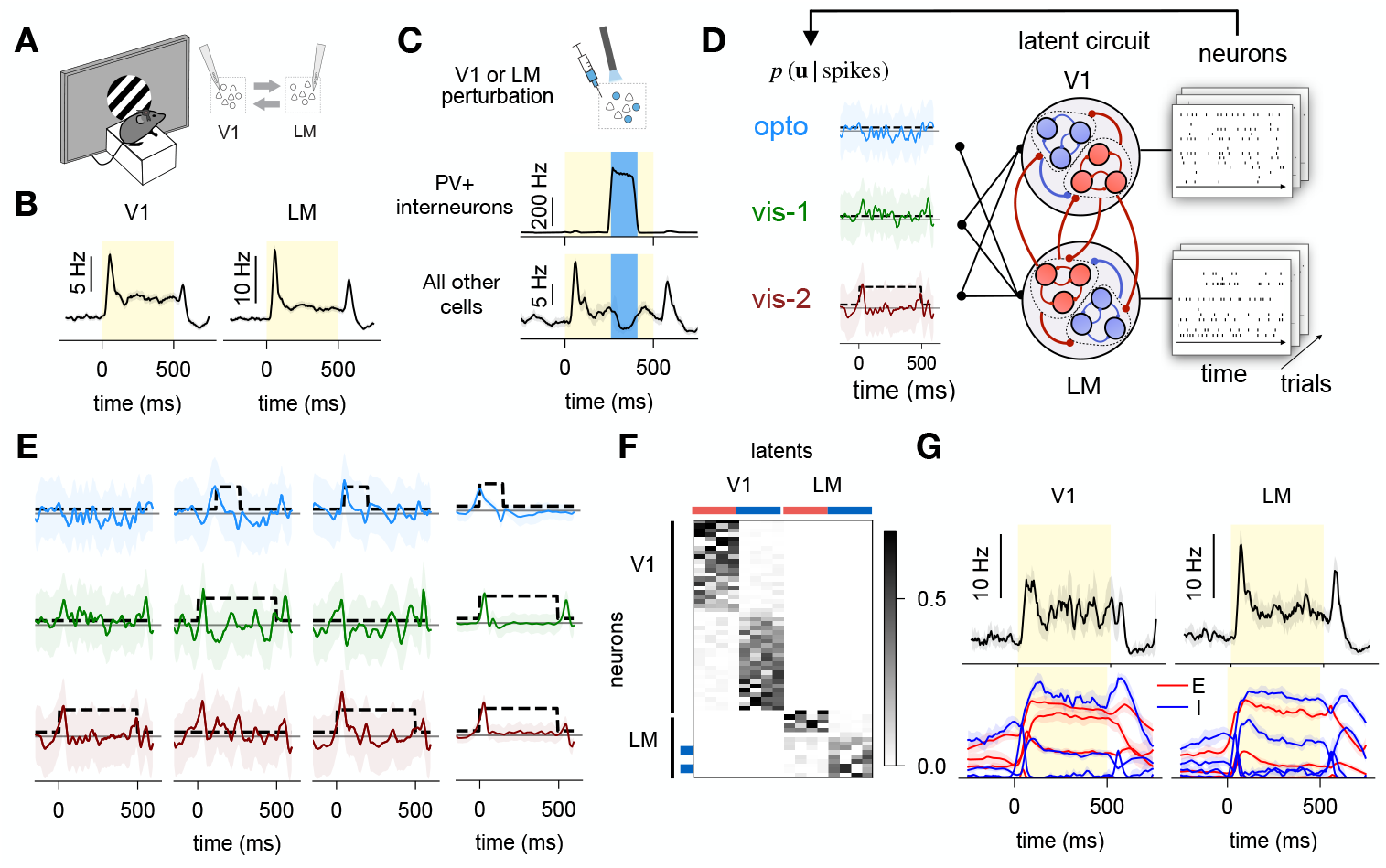
Modeling input-driven dynamics in the V1-LM network during visual processing. **(A)** Head-fixed, stationary mice were presented with two differently oriented stationary grating stimuli (45° or -45°), only one of which was rewarded. Mice reported the rewarded stimulus by licking a spout, which triggered the delivery of the reward (go/no-go). Paired neural recordings were performed in retinotopically matched regions of V1 and LM with silicon probes in PV-Cre mice. **(B)** Trial- and neuron-averaged spiking activity in no-go trials in V1 (left) and LM (right). A total of 194 neurons in V1 and 228 neurons in LM were recorded in 7 mice across a total of 513 ±110 (mean ± std) correct trials per mouse across two stimuli. **(C)** Top: In some trials, either V1 or LM was silenced through light-mediated activation of parvalbumin-expressing inhibitory cells expressing ChR2. The light onset was randomly chosen in each trial amongst 8 different times, spanning the duration of the stimulus uniformly (at 65 ms intervals), with a total of 449 ± 98 (mean ± std) silencing trials per mouse. Bottom: Neuron- and trial-averaged spiking activity of the optogenetically stimulated PV+ neurons (top) and all other neurons (bottom) in an example animal, for one laser delay. Biologically-constrained latent circuit model of V1-LM, with dynamics driven by 3 external inputs whose time course is inferred on a single trial basis. Dashed lines indicate the time-varying standard deviations of the (zero-mean) prior distributions over these inputs. Solid lines and shaded areas indicate posterior mean and standard deviation respectively, in one example trial, estimated from 100 posterior samples and smoothed with a running average of 25ms for visualization. **(E)** Left: Inferred time course of inputs for three example trials. Each column shows three input channels for one trial (optogenetic perturbation in blue, go stimulus in green, and no-go stimulus in red). The prior standard deviation (dashed line) indicates the presence of each input in the trial: no-go stimulus in the first trial, go stimulus paired with optogenetic perturbation in the second trial, and no-go stimulus paired with optogenetic perturbation in the third trial. Shaded area is the posterior standard deviation. Right: Trial-averaged time course of the three input channels, aligned to the input onset, shown as mean and standard deviation (shaded area) of the posterior mean across all trials. For each input channels, trial averages were calculated from trials where that input was present. All traces were smoothed with a running average of 25ms for visualization. **(F)** Example readout matrix (***C*** in Equation 2) in the fitted model, depicting the mapping from the latent units (columns, divided into two areas, and into excitatory (red) / inhibitory (blue) subpopulations within each area) to the recorded neurons (rows; blue bars mark PV cells identified by optogenetic perturbations in LM). **(G)** Top: Average recorded activity in V1 (left) and LM (right) during the no-go visual stimulus in an example animal. Bottom: Corresponding activity of the excitatory (red) and inhibitory (blue) latent units. Shaded areas around mean traces in B, C, and G denote 95% confidence intervals (±2 s.e.m.).

We built circuit models that explicitly incorporated known aspects of cortical circuit organization, in particular the excitatory nature of long-range connections between areas and local excitation-inhibition dynamics. In these models, the time course of spiking activity in V1 and LM was explained by the recurrent dynamics of the latent circuit (Figure 1D). These dynamics were driven by time-varying inputs that we inferred for each trial, reflecting any unobserved signals external to the V1-LM circuit such as sensory or optogenetic stimuli.

Specifically, the latent circuit’s activity ***z***(*t*) evolved according to

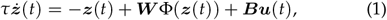

where *τ* = 20 ms is the characteristic neuronal membrane time constant, ***W*** is the latent circuit connectivity, Φ(·) is a soft-rectified nonlinear activation function, and ***u*** is a set of trial-specific external input signals that enter the dynamics through the input matrix ***B*** (Methods). The activity of this latent circuit was used to describe firing rate fluctuations in the observed V1 and LM neurons, according to

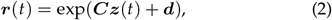

where ***C*** is a readout matrix specifying the way in which each recorded neuron relates to the latent units, and ***d*** is a vector of constant offsets. Action potentials were modelled as Poisson processes given these time-varying firing rates.

The latent circuit was partitioned into two areas, which mapped onto V1 and LM neurons respectively. Moreover, the recurrent connectivity matrix ***W*** was constrained so that each area was composed of separate populations of excitatory and inhibitory units, and long-range connections between the two areas originated exclusively from the excitatory units (Methods). Although we did not know the E/I identities of most of the recorded neurons, we used a specific sparsity penalty on ***C*** to discourage any nonsensical, simultaneous association of a neuron with both E and I latents subpopulations (Methods). This soft constraint encouraged the model to learn to label each neuron as *either* E *or* I.

To fit the model, we used iLQR-VAE [Schimel et al., 2022], a method ideally suited to learning the dynamics of a circuit when the detailed time course of external inputs is unknown and must therefore be inferred in each trial. Importantly, here we did have some knowledge of *what* input signals might have driven the circuit in a given condition and *when*. iLQR-VAE lets us incorporate such information in the form of condition-specific, time-varying statistical priors over the input ***u***(*t*) in Equation 1. Thus, we used three input channels reflecting the two visual stimuli and the optogenetic perturbation events. The mapping from inputs to latents, ***B***, was constrained such that the input channel with the optogenetic perturbation prior could only target the inhibitory latents of the stimulated area for each animal. For each channel, we learned two prior variances: the higher variance was used during the time the corresponding stimulus was on, and the lower one outside those epochs (Figure 1D, dashed lines). This encouraged the model to use larger inputs when the stimuli were present, while retaining flexibility with respect to their exact time course. iLQR-VAE then inferred this time course on a single trial basis, by computing a posterior distribution over the input signals in each channel conditioned on the observed neural data (Figure 1D and E, solid lines).

The parameters of the model (***W***, ***B, C, d*** and the prior variances) were obtained by maximizing the likelihood of the observed spike trains. For each animal, we performed multiple fits starting from random initializations (Methods), and found that each fit robustly attributed a definite E or I identity to each observed neuron (Figure 1F). For most fits (78.24%), the model correctly labelled all of the directly photo-stimulated neurons (known to be PV+ inhibitory cells) as inhibitory (Figure 1F, cyan mark); we rejected the few models where these cells were mislabelled. Finally, for each animal we selected the model with the best goodness of fit on held-out data (see below, and Methods).

## The model captures trial-by-trial variability

We first characterized how well the learned models captured single trial activity in our recorded neurons. For each trial, we could leave one neuron out, and let the model infer the time course of its firing rate given the activity of the other neurons (Figure 2A). Based on this single-trial firing rate, the model then attributed a (Poisson) likelihood to each spike for that neuron. On average, this single-trial likelihood was greater than that predicted by the PSTH of the same cell ob-tained by averaging over the other trials in the same condition (Figure 2B, ‘residual likelihood’). In other words, our latent circuits captured the spatio-temporal structure of our recordings beyond condition averages. Accordingly, our models also captured the structure of pairwise covariances in neural activity (Figure S3). Importantly, the models did not significantly suffer from the circuit constraints we imposed; they explained the single-trial data just as well as fully unconstrained models (Figure 2A-B, gray; Methods).

**Figure 2.**
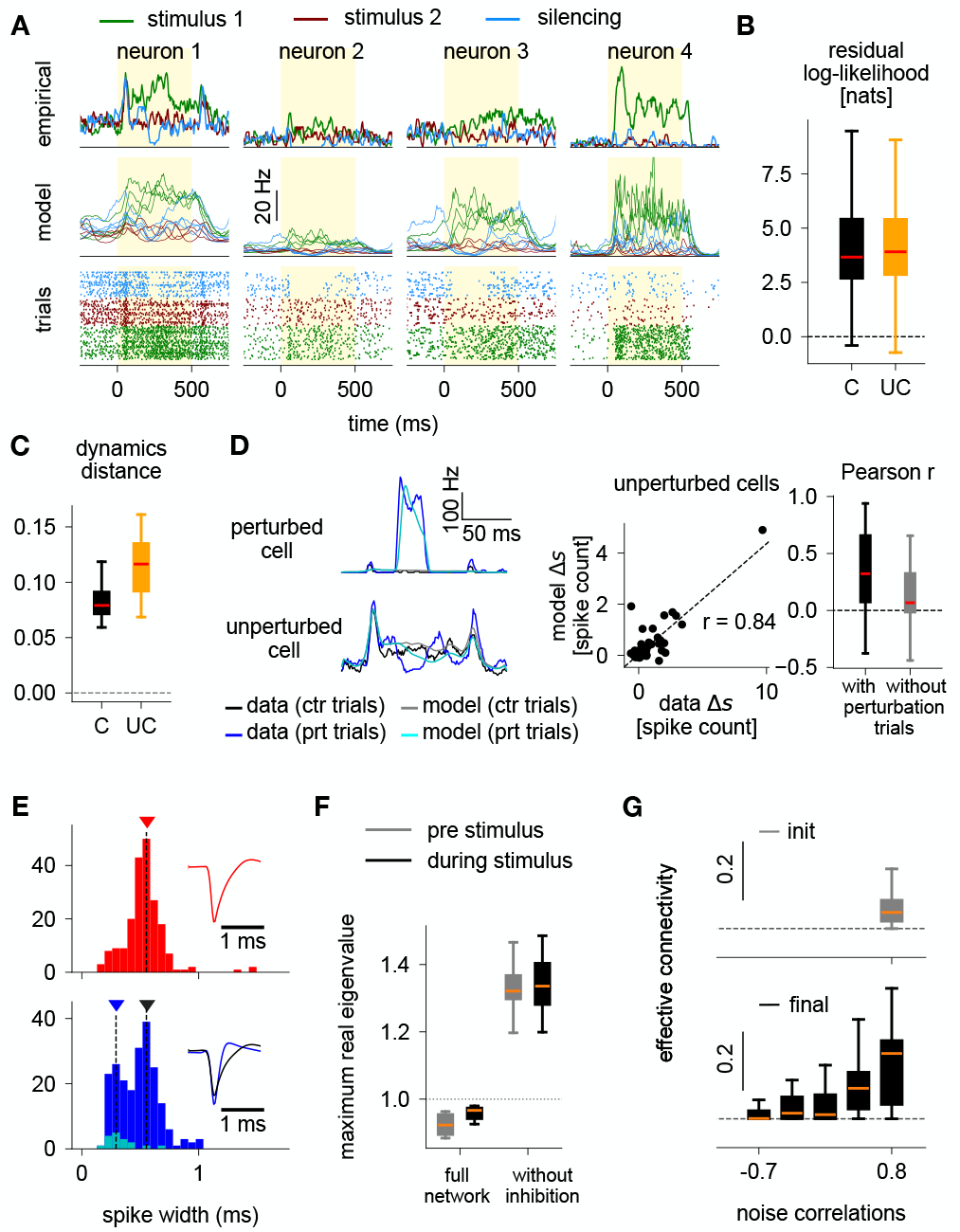
Circuit-constrained models capture the statistics and underlying mechanisms of neural activity. **(A)**Top: trial-averaged empirical firing rates, smoothed with a running average of 25 ms, for 4 example neurons in one animal. Different colors denote the two visual stimuli (red and green) and one silencing condition (cyan). Middle: corresponding model-predicted firing rates in individual trials (4 trials per condition), given the spikes observed concurrently in all other neurons. Bottom: corresponding spike rasters in the same three conditions, for the same four neurons. **(B)** Residual log-likelihood of the predicted firing rates of active cells (test trials, see Methods) for the constrained (‘C’, black) and unconstrained (‘UC’, orange) models. *n* = 246 neurons across 7 animals, ‘C’ vs. ‘UC’ paired p-value = 6 × 10^−7^, unpaired p-value = 0.23. **(C)** Between-animal similarity in model dynamics, linearized around the stimulus-period activity. Similarity is computed as an average pairwise Procrustes distance (see Methods), calculated separately for constrained (‘C’) and unconstrained (‘UC’) models (paired p-value = 6.7 × 10^−6^). **(D)** Left: trial-averaged activity of two example neurons, obtained either from the data (control no-go trials in black and one perturbation condition in blue), or by running the learned dynamics forward given condition-specific inferred inputs and artificial perturbations (see Methods; control no-go trials in gray and one perturbation condition in cyan). The top cell is directly perturbed by the optogenetic perturbation, while the bottom cell is only indirectly affected. Center: change of neural activity (relative to control trials) predicted by the model in response to a simulated optogenetic perturbation, as a function of the experimentally observed response difference between laser and corresponding control trials. These are shown for all cells in V1 and LM that were not directly perturbed, and in one animal. *r* denotes the Pearson correlation coefficient between the predicted and true change (see Methods and Figure S4). Right: distribution of Pearson correlation coefficients (c.f. middle) across animals, for models trained with perturbation data (black) and without perturbation data (gray) (Paired *p* = 0.0003). **(E)** Histograms of spike widths for the neurons labeled by the model as excitatory (top, red) and inhibitory (bottom, blue; known PV cells shown in cyan). Insets show the average spike waveforms (± 2 sem) for those neurons lying around the marked peaks. **(F)** Distribution across animals of the maximum real part of the eigenvalues of the latent circuit dynamics, linearized either before (gray) or during (black) stimulus presentation (pooled across both go and no-go trials). This is shown for the full model (left), and in the absence of inhibition (right; see Methods). **(G)** Effective connectivity (see Methods) plotted against noise correlations in control no-go trials, for pairs of latent circuit units. This is shown at model initialization (top gray), and after training (bottom black).

## The model infers circuit dynamics that are consistent across animals and captures key aspects of V1-LM cortical physiology

That constrained and unconstrained models explain the data equally well, despite being entirely different families of dynamical systems, raises a concern: have our constrained models learned dynamics that really capture the mechanics of the underlying V1-LM circuit?

A first indication of faithful dynamics reconstruction is the consistency of the learned solutions across animals. We evaluated the distance between the inferred latent flow fields between pairs of animals, accounting for an arbitrary rotation of the latent state space for each animal (Methods). This analysis revealed that the dynamics learned by our constrained latent circuits were broadly consistent across animals – indeed more consistent than in unconstrained models (Figure 2C).

As a second, stronger test of accurate dynamics reconstruction, we probed the responses of our models to *internal* perturbations of the inhibitory cells, and compared those to the responses observed in the V1-LM data during optogenetic perturbations. Specifically, we simulated single perturbation trials by directly providing positive input to the inhibitory latent units of the relevant area for the entire duration of the photo-stimulation, whilst replaying the external inputs inferred on a control trial (no photo-stimulation) for the relevant condition (go vs. no-go). We adjusted the amplitude of each perturbation input (four parameters per animal) in order to match the trial-averaged norm of the responses of the known PV cells in the stimulated area. We then evaluated the model-predicted change in firing rates in the other neurons (per-condition averages). These predictions were positively correlated with the corresponding firing rate changes observed in the data (Figure 2D; average Pearson *ρ* = 0.34). In contrast, models trained using only no-perturbation trials failed to capture the sensitivity of the V1-LM circuit to photoinhibition (Figure 2D, gray; average Pearson *ρ* = 0.12), highlighting the importance of optogenetic manipulations for accurate neural system identification.

As a third indication that our model has learned the correct circuit structure, we looked at the excitatory/inhibitory identity that it assigned to each neuron in our recordings. Whilst we used the assigned identity of the known PV cells in the photostimulated area as a criterion for model selection, we could study the identity assigned to the other recorded neurons. Inhibitory neurons in the cortex are known to exhibit a bimodal distribution of spike widths: fast-spiking (PV) interneurons exhibit narrow spike waveforms, whilst other (non-PV) neurons have slower action potentials similar to that of excitatory neurons [Rudy et al., 2011]. Consistent with this known aspect of cortical electrophysiology, we found that the neurons which the model deemed inhibitory had a bimodal distribution of spike widths (Figure 2E). The mode of the histogram corresponding to broader spikes aligned well with the distribution of spike widths in the neurons classified as excitatory by the model.

Finally, the dynamics inferred by the model are consistent with previous studies of the mammalian visual cortex. In particular, the network operates in the inhibition-stabilized regime (Figure 2F; Ozeki et al., 2009, Ahmadian and Miller, 2021), whereby the excitatory subnetwork is unstable on its own but stabilized by feedback inhibition. In fact, the network is inhibition-stabilized even in the absence of visual stimulation, as previously shown in mouse V1 [Sanzeni et al., 2020]. Moreover, noise correlations in the latent circuit reflect the strength of excitatory connectivity between pairs of latent units (Figure 2G), as observed in mouse visual cortex [Ko et al., 2011]. Importantly, this relationship was not present at initialization, but arose after fitting the model to the data.

## Contribution of external and recurrent inputs in shaping cortical visual responses

Having established the validity of our model fits, we then used the resulting latent circuits to dissect the roles of various structural components of the V1-LM network in shaping its sensory responses. To do this, we focused on several key features of the learned latent circuit connectivity, systematically and individually down-modulated their strengths, and quantified the effect of these modulations on the circuit’s responses to sensory stimuli. Only no-go trials, with no reward or licking-related movement, were used for this analysis (Figure 3)

**Figure 3.**
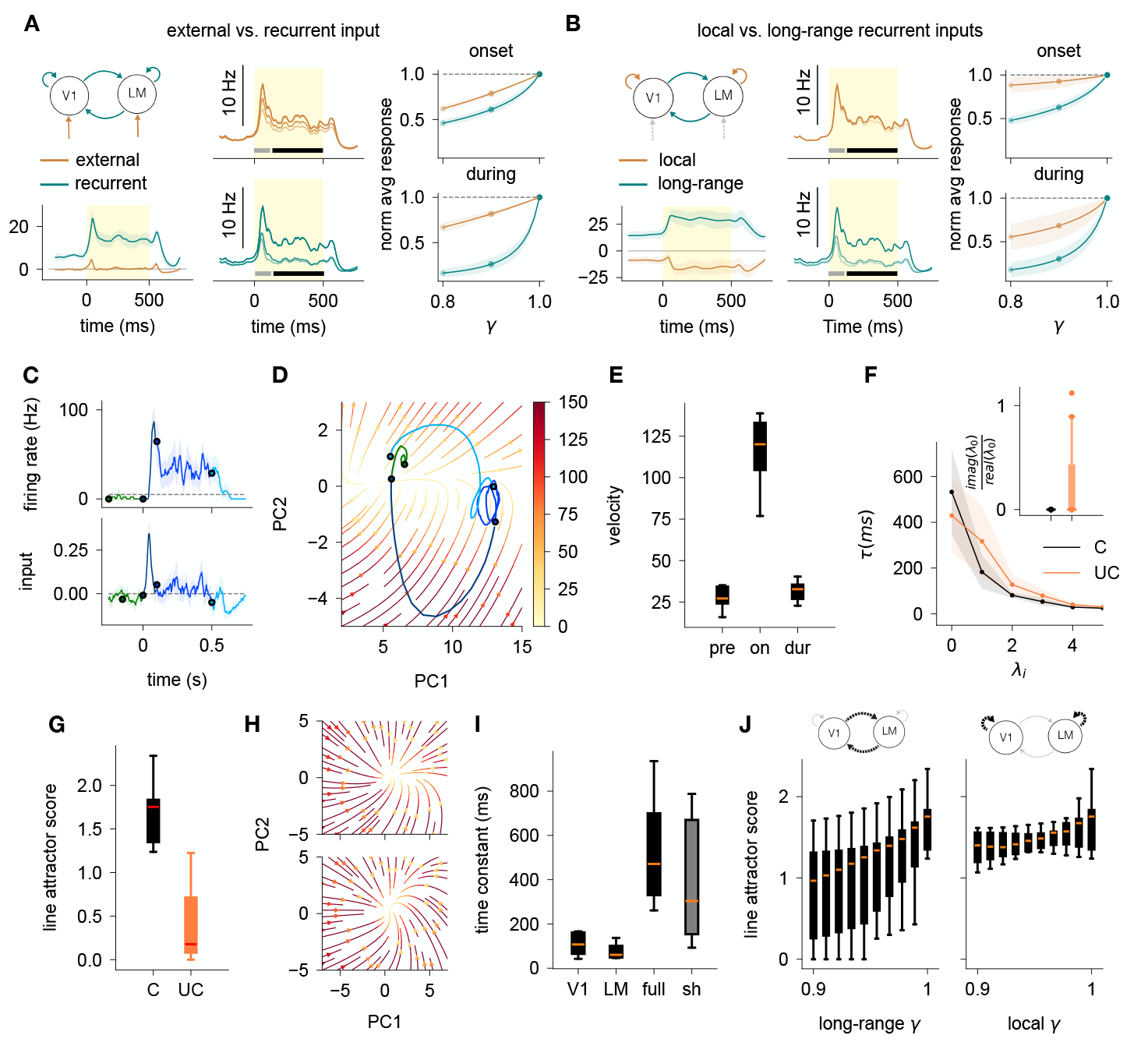
Maintenance of activity via a slow mode emerging from interacting E/I networks. **(A)**Left top: Schematic showing the external (teal) and recurrent (burnt orange) inputs in the V1-LM circuit. Left bottom : Average external and recurrent inputs, in no-go trials across all animals (shaded area denotes 2 sem). Middle top : Average network activity in no-go trials in an example animal, as we scale down the gain of the external inputs (see Methods). We use gain values of 0.8, 0.9 and 1, ordered from light to dark. The grey and black bars denote the stimulus onset and during the stimulus. Middle bottom : Same as top, for recurrent inputs. Right : Sensitivity, i.e change in the response of the network (see Methods) as we scale down (by *γ*) the external (burnt orange) or recurrent (teal) inputs. This is shown at stimulus onset (top; first 100ms of the stimulus) and during the stimulus presentation (bottom; 100-500 ms after stimulus onset). **(B)** Same as (A), but comparing local and long-range recurrent inputs (see Methods). **(C)** Trial-averaged firing rate of an example neuron during no-go trials, smoothed at 25 ms (top) and the trial-averaged inferred external input to the circuit, averaged over all latent units (bottom). The colors indicate different time segments. **(D)** Flow field of the dynamics for the same animal as (C), projected in the subspace defined by the top 2 PCs of the latent activity (see Methods). The color bar indicates the magnitude of the velocity. The trajectory represents the projection of the trial-averaged inferred latent activity in no-go trials, with time segments color-coded as in (C). **(E)** Velocity of the autonomous latent dynamics (i.e latent dynamics in the absence of external input), averaged either over the pre-stimulus period (−400-0 ms), around stimulus onset (0-100 ms), or during the stimulus (100-500 ms). **(F)** Relaxation time constants (mean ± 2 sem across all animals; see Methods) of the linearized dynamics in the 100-500 ms time window of no-go trials. This is shown for constrained models in black and unconstrained models in orange. The inset shows the absolute value of the imaginary to real ratio of the eigenvalues corresponding to the slowest direction. **(G)** Distribution across animals of the line attractor score (see Methods) of the dynamics, linearized around the mean activity in the 100-500 ms time window of no-go trials, for the constrained (black) and unconstrained (orange) models. Flow field of the V1-only (top) or LM-only (bottom) dynamics, for the same example animal and trials as in (D), projected in the subspace defined by the top 2 PCs of the latent trajectories in each area (see Methods). **(I)** Distribution across animals of the lowest relaxation time constant of the dynamics in the full networks (constrained models) and the V1-only or LM-only networks. The gray box corresponds to the distribution of slowest time constants in networks of the size of a single area, randomly sub-selected from the full networks. **(J)** Line attractor score of the V1-LM network, as we scale down the long-range (left) or within-area (right) connections by a factor *γ*.

We began by dissociating external and recurrent inputs to the latent circuit. Whilst the average external input to each neuron was mostly transient, i.e. confined to the onset and offset of the sensory stimulus, the corresponding recurrent input remained elevated for the whole stimulus duration (Figure 3A, left), mirroring the period of sustained activity across two areas during the stimulus epoch (recall Figure 1G). Even modest down-scaling of all recurrent weights during the stimulus (Figure 3A, center bottom, black bar) could nearly abolish these sustained responses. Similar down-scaling of recurrent connectivity during stimulus onset (Figure 3A, center bottom, gray bar) had a weaker effect (Figure 3A, right; compare top and bottom green curves). Modulation of the external input weights had a weaker effect still (Figure 3A, center and right), indicating that sustained activity arose primarily from recurrent connections, with external inputs triggering the onset response.

Next, we characterized the differential contributions of local vs. long-range connections. While the net local inputs were smaller and negative (inhibition-dominated; Haider et al., 2013), the long-range inputs were stronger (and positive by design; Figure 3B, left) leading to positive net recurrent inputs. Moreover, modulating local and long-range connection strengths separately revealed that stimulus-epoch sustained activity depended strongly on long-range interactions, but more weakly on within-area interactions. Together, these sensitivity analyses suggest a mechanism for sustained sensory responses in the V1-LM circuit that relies on across-area reverberation of activity, mediated by bidirectional long-range connections.

## Sustained sensory responses are maintained by approximate line attractor dynamics across V1 and LM

To further characterize the origin and properties of V1-LM reverberation induced by transient inputs (Figure 3C), we analyzed the activity flow field in the latent circuits. In the subspace defined by the two principal components of latent activity, the autonomous flow of the latent circuit’s dynamics (i.e. latent trajectories obtained in the absence of external inputs) primarily converged towards a line of slow dynamics (Figure 3D, one example mouse). Consistently across mice, the latent state trajectories underlying the neural data spent most of the stimulus epoch near this line of weak flow, only briefly leaving this region at stimulus onset and offset in response to transient external inputs (Figure 3E). This picture is highly suggestive of line attractor dynamics [Ganguli et al., 2008, Mante et al., 2013, Nair et al., 2023, Sylwestrak et al., 2022], a regime characterized by slow decay of activity along a select direction in state space, with all other directions decaying more rapidly. Mathematical analysis of the time constants present in the latent circuits (Figure 3F, Methods) revealed such a gap, with local dynamics around the stimulus-evoked response largely dominated by a slow mode with a timescale of ∼ 400 ms, which is 20 times longer than the characteristic time constant of single neurons in our model (20 ms). Although the second longest time constant was also slow (≫20 ms) – a point we will return to below (Figure 4) – it was significantly shorter than the slowest, reflected in a high “line attractor score” (Figure 3G; Methods).

**Figure 4.**
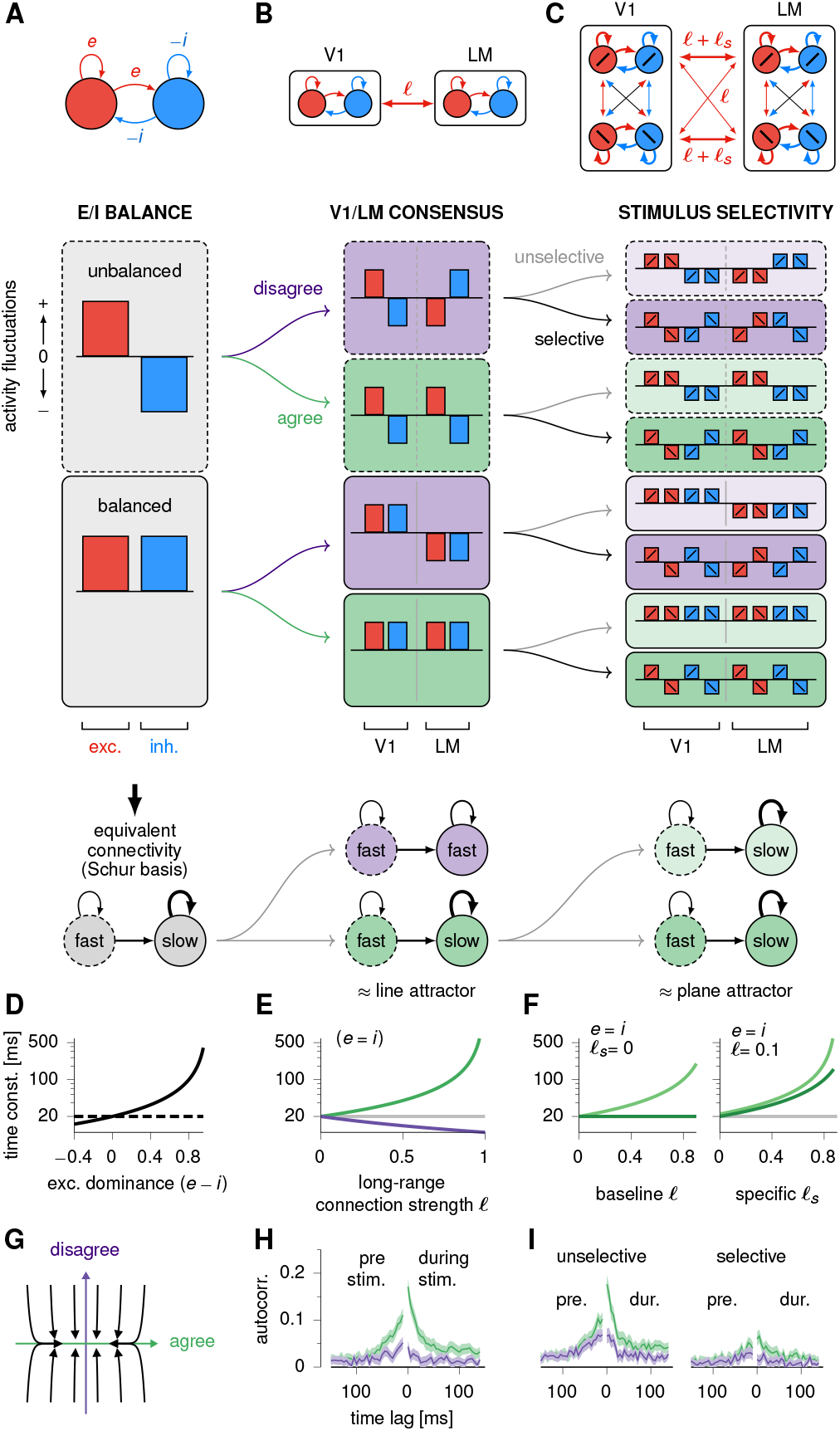
Network time constants in minimal models of interconnected E/I circuits. **(A)**In a canonical E-I circuit (top), momentary activity can always be expressed as a linear combination of two modes (middle): an “unbalanced” mode (dashed outline) and a “balanced” mode (solid outline). Recurrent E-I connectivity is equivalent to feedforward connectivity from the unbalanced to the balanced mode (bottom; Murphy and Miller, 2009). The unbalanced mode exhibits fast dynamics, whereas the balanced mode can evolve more slowly depending on the degree of excitatory dominance (c.f. D). **(B)** Minimal E-I model of V1 and LM (top), where connectivity within each area is of the same form as in (A), and each area excites the other via long-range connections of strength 𝓁. In this model, activity can be decomposed into four modes (middle): a pair of balanced & unbalanced modes in which V1 and LM activities are anti-aligned (purple, ‘disagree’), and a similar pair in which they align (green, ‘agree’). These two pairs of modes are decoupled, and interactions within each pair are effectively feedforward (bottom). The dynamics are slowest along the mode of balanced agreement (c.f. E), resulting in an approximate line attractor. **(C)** This minimal model can be extended to accommodate selectivity to ± 45° visual gratings, by splitting each E/I population into two differentially selective subpopulations. All connection types (local E, local I, long-range E) are composed of an unselective baseline and a selective (like-to-like) components (top). This connectivity structure gives rise to two versions (unselective, pale / selective, dark) of each of the four modes in B (middle), and results in slow dynamics in the two modes of balanced agreement (approximate ‘plane attractor’). **(D-F)** Time constants of the balanced mode(s) for each model (colors as in A-C), as a function of key connectivity parameters. In (F), only the two slow modes are shown. **(G)** Flow field of the dynamics of the model in (B) in the activity plane spanned by the two balanced modes, showing convergence onto the ‘agree’ mode. Each line is obtained by integrating the network’s dynamics starting from a different initial condition in that plane. **(H)** Autocorrelation function of the neural data pre-(left half) and during stimulus (right half) projected onto the ‘agree (resp. disagree) balanced’ modes (green resp. purple). These two modes correspond to the sum (resp. difference) of the average V1 and LM spiking activities. Traces are mean ± 95% confidence intervals. pre: Δ_max_ = 0.06, *p <* 10^−5^. during: Δ_max_ = 0.13, *p <* 10^−5^. **(I)** Same as (H), but for neural activity projected onto the pair of unselective agree/disagree modes (left) and analogous selective versions (right) defined in the main text. unselective, pre: Δ_max_ = 0.035, *p* = 0.0006. unselective, during: Δ_max_ = 0.11, *p <* 10^−5^. selective, pre: Δ_max_ = 0.022, *p* = 0.003. selective, during: Δ_max_ = 0.054, *p <* 10^−5^.

Importantly, the approximate line attractor we identified in the latent models arose from the constraints we imposed on the structure of the circuit. Indeed, whilst unconstrained model fits did also produce slow dynamics (Figure 3G, orange), they exhibited less consistent line attractor scores, primarily because their slowest modes were occasionally oscillatory (and therefore planar; inset). Moreover, we found that the line attractor arose specifically from the long-range excitatory interactions between V1 and LM. First, the line attractor score was sensitive to modulation of the long range, but not the local, connections (Figure 3J). Second, each area considered separately (i.e. with long-range connections removed) did not exhibit any line attractor (Figure 3H) and had substantially faster dynamics (Figure 3I). Although weaker reverberation of activity in those isolated areas could in principle reflect their smaller sizes, randomly thinning both latent sub-circuits by eliminating half of their units did yield significantly slower dynamics than those of the isolated areas (Figure 3I, gray).

## A minimal model of the V1-LM circuit explains the emergence of a line attractor

To understand the circuit mechanisms that underlie the emergence of a line attractor across V1 and LM, we considered simplified models of multi-area excitation/inhibition (E-I) networks. As a starting point, we recall a canonical model of cortical E-I circuits, with one E and one I population recurrently connected as shown in Figure 4A (top), with E-I weight parameters *e* and *i*. In these networks, activity can be generically decomposed into two main motifs (Figure 4A, middle): E-I imbalance (with the E population firing more than average, and the I population firing less; dashed boxes) and E-I balance (both populations firing in the same way; solid boxes). In this modal decomposition, the recurrent connectivity is more easily interpreted: it acts to transiently amplify any momentary imbalance in network activity into balanced activity (Figure 4A, “Schur basis”; Murphy and Miller, 2009). Whilst E-I imbalance is typically short-lived, balanced activity may linger depending on the level of excitatory dominance in the recurrent connectivity (Figure 4A, bottom; Supplementary Material S2).

Next, we extended the canonical single-area E/I model to two interacting areas, yielding an idealized reduction of our latent circuit models of V1 and LM. In this model, each area is modelled as an E-I circuit as above, and they interact via long-range excitatory connections of strength 𝓁 (Figure 4B). Mathematical analysis of this model revealed a similar kind of feedforward connectivity as for single-area E-I networks, now for two different sets of unbalanced/balanced modes. In the first set, the two areas fluctuate congruently such that their patterns of E-I activities – whether balanced or unbalanced – are aligned (“agree”, green boxes). In the other set, these patterns are anti-aligned across the two areas (“disagree”, purple boxes). Recurrent connectivity now acts separately on each set, with transient amplification of congruent/incongruent E-I imbalance into the corresponding balanced pattern. Notably, long-range excitatory connectivity has an opposite effect on each set of modes: it acts to slow down activity where the two areas agree, and speed up the decay of any disagreement (Figure 4E; Supplementary Material S2). This separation of timescales gives rise to approximate line attractor dynamics in the combined circuit (Figure 4G), as observed in the latent circuit models we had obtained from data (recall Figure 3D-G). Moreover, the model clarifies that the line attractor arises specifically from long-range connections as previously shown in Figure 3H-J. In addition, the model confirms that the line attractor score (which depends directly on the timescale separation) should grow with the strength of those long-range connections as in Figure 3J.

Notably, this simplified model of V1-LM interactions not only provided a qualitative explanation for the emergence of a line attractor, it also matched the dynamics of our latent circuit models quantitatively. Indeed, with optimally chosen parameters, this 4-dimensional network could account for 65% of the impulse response of the (linearized) 16-dimensional network (Figure S8).

Importantly, the simplified model of V1-LM interactions out-lined above makes a prediction that can be tested independently of our latent circuit model fits. Specifically, V1-LM activity projected along the balanced-agree mode should exhibit slower fluctuations than along the balanced-disagree mode. To verify this prediction experimentally without relying on the latent circuit model, we estimated the contribution of each of these two modes to the momentary activity of the recorded neurons in our dataset. This was done by separately averaging the activity of V1 and LM neurons to estimate local balance in each area, and then taking the sum (agree) and the difference (disagree) of these local averages. As predicted, we found that the empirical ‘balanced-agree’ mode had a longer autocorrelation decay time than its ‘disagree’ counterpart (Figure 4H; Methods), both before (left) and during (right) the presentation of the sensory stimulus.

More generally, the model predicts slower dynamics along the balanced-agree mode compared to *any* other mode, including the unbalanced modes. Testing this more general prediction without referring to our latent circuit models is difficult, because estimating momentary E-I imbalance in V1 or LM directly from the neural data requires knowing the E-I identities of all cells. Nevertheless, identifying these modes based on model-predicted cell identities (c.f. Figures 1 to 3) allowed us to confirm this more general prediction (Figure S7A).

## Multi-area consensus on stimulus presence and identity via selective long-range interactions

The selective slowing down of activity patterns where V1 and LM “agree”, and concurrent quenching of patterns where they disagree, can be seen as a circuit mechanism for consensus building (Figure 4G). We wondered about the generality of this mechanism: whilst the minimal 2-area model of Figure 4B gives rise to consensus regarding whether or not a stimulus is present, a similar mechanism could also underlie consensus about stimulus identity. We hypothesized that this second mode of consensus might also account for the second slowest mode in the learned dynamics (Figure 3F), which the simple reduced model introduced above was unable to explain.

To explore this hypothesis, we took a similar modelling approach as above. We constructed a more detailed reduced model of a 2-area network (Figure 4C) that incorporates feature specificity in its connectivity (see Supplementary Material S2). Each area was split into two E-I sub-circuits that were differentially driven by two orthogonally oriented stimuli (corresponding to go and no-go stimuli in our experiments). Recurrent E-I connectivity in each area had a degree of specificity that we could vary, i.e. connectivity could be made stronger within, compared to between, the two local sub-circuits with different stimulus preference. Similarly, long range excitatory connection strengths included both a baseline (𝓁_0_) and a specific (𝓁_s_) component (Methods).

The effect of the connectivity on the dynamics of this circuit could again be understood by considering a modal decomposition similar to Figure 4B, which included (i) patterns of E-I imbalance/balance, in which (ii) the two areas could either agree (green) or disagree (purple), and which (iii) were either stimulus selective (dark) or unselective (light). We found that this circuit would predominantly dwell in two of these activity modes: the ‘balanced-agree-selective’ mode, and the ‘balanced-agree-unselective’ mode, both of which were characterized by long time constants. As before, all ‘unbalanced’ and ‘disagree’ modes were associated with comparatively faster decay times. The slow decay of the ‘balanced-agree-unselective’ mode relied on strong long-range connections regardless of specificity (𝓁_0_ or 𝓁_s_; Figure 4F, left), whilst the slow decay of the ‘balanced-agree-selective’ mode required strong *specific* long-range connections (𝓁_*s*_; Figure 4F, right). Thus, this circuit supports the dynamic formation of a consensus across V1 and LM about both the presence of a stimulus and its identity. The presence of a second slow mode made this 8D reduced model an even better quantitative match to the linearized dynamics of the full 16D model (Figure S9; 77.3% of variance captured in the impulse response). Additionally, we also verified that the two slowest modes in the dynamics of our latent circuit models aligned well with the unselective and selective balanced-agree modes (Figure S7F).

We could again articulate the model’s predictions regarding the relative timescales of these different activity modes, and use our neural recordings to test these predictions. In particular, the model predicted that the network’s activity should fluctuate slower along the two main modes of consensus than along the corresponding modes of disagreement. To test this hypothesis independently of our model fits, we estimated the degree of engagement of each neuron in the four ‘agree/disagree-selective/unselective’ modes based on its observed responses, and assessed the slowness of population activity projected onto these modes. Specifically, we first extracted the sensitivity of each recorded population (V1 or LM) to the presence of a stimulus irrespective of its identity by taking the difference of its population activity vector after and before stimulus onset, denoted by 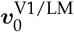 (Methods). We then defined the ‘agree-unselective’ mode as 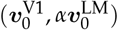, and the ‘disagree-unselective’ mode as 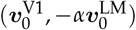, where *α* is a scaling factor that accounts for unequal sampling of V1 vs. LM neurons in our recordings (Methods). Similarly, by computing the differences 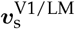 between population responses to the go and no-go stimuli, we could define the ‘agree-selective’ and ‘disagree-selective’ modes as 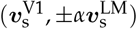 respectively. We then projected the activity of all recorded neurons onto these four modes and computed the autocorrelations of the resulting signals (Figure 4I). As predicted by the model, activity fluctuated slower along both ‘agree’ modes, compared to the corresponding ‘disagree’ modes. This was true both for activity taken before and during the stimulus presentation. The relative slowness of the ‘agree-selective’ mode thus suggests some degree of specificity in the long-range connections between V1 and LM, consistent with experimental findings [Ding et al., 2023].

## Functional consequences of consensual dynamics

To explore the functional significance of slow unselective and selective consensual dynamics across V1 and LM, we revisited our minimal selective model (Figure 5A, top), and examined its dynamics along the agree/disagree modes identified earlier. Specifically, we provided the model with a stimulus whose time course captured both the strong transient and weaker sustained characteristics of the input which we had inferred from the recorded spiking data (Figure 5A, middle). By varying the degree to which the stimulus drove (i) each area, as well as (ii) each sub-population therein, we could manipulate the degree of ‘input agreement’ between V1 and LM about (i) the presence of a visual stimulus and (ii) whether it is oriented at +45 or − 45 degrees (Figure 5B and E, gray insets). We could then examine any emergent consensus in the network’s response. For example, the stimulus pattern shown in Figure 5B (gray) drives V1 and LM in opposite directions, increasing V1 activity while suppressing LM (evidence for the presence of the stimulus in V1, and against it in LM). Mathematically, this stimulus recruits both the agree- and disagree-unspecific modes, leading to input ambiguity. In the absence of long-range connections between V1 and LM, the network’s response directly reflects this lack of consensus in the input (Figure 5B, left; C and D, black). However, with increasingly strong long-range connections, the network selectively amplifies the input contribution to the agree-unselective mode, while suppressing it for the disagree-unselective mode, thus allowing an inter-area consensus to dynamically emerge on the presence of a stimulus (Figure 5B-D, orange). Importantly, this consensus is contingent on bidirectional inter-area reverberation of activity, and is significantly diminished if the feedback connections from LM to V1 are ablated (Figure S10).

**Figure 5.**
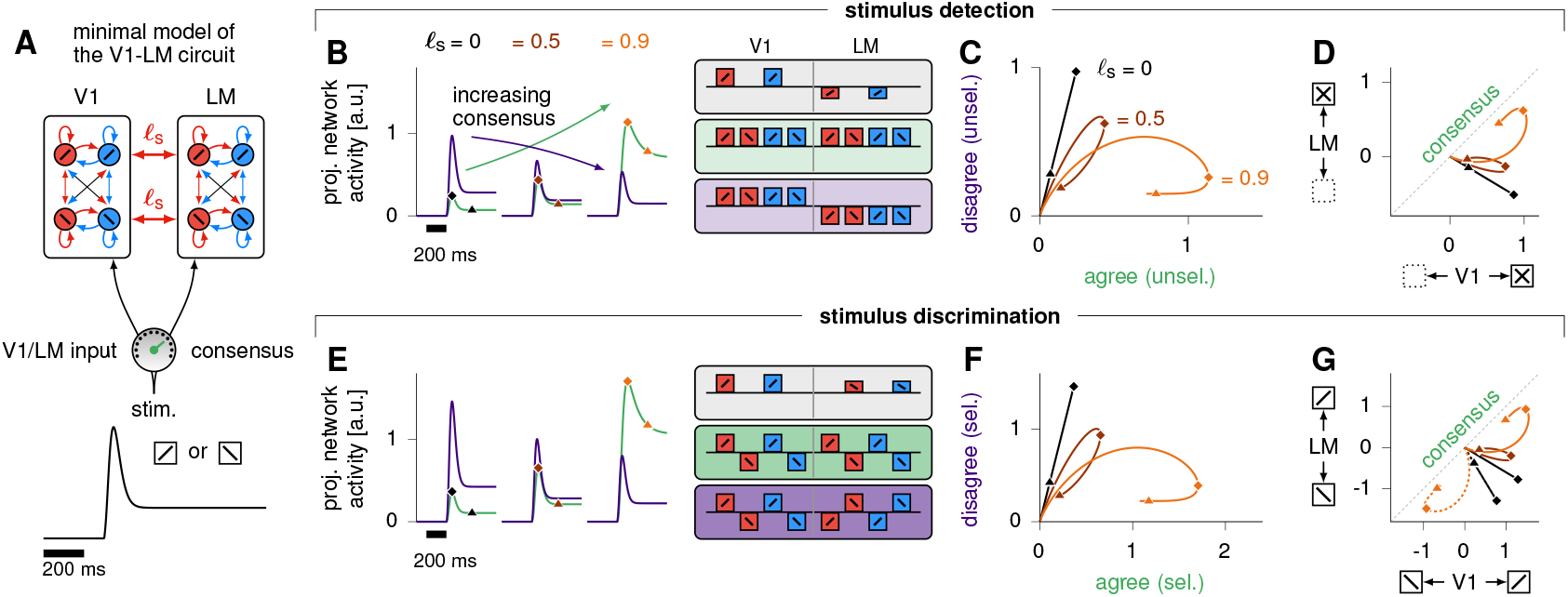
Dynamic emergence of consensus via long-range connections in minimal models of interconnected E/I networks. **(A)** Schematics of the minimal selective model of the V1-LM circuit, driven by an external stimulus that can target either one of the two E/I pairs in each area depending on their orientation preference (⊘ vs.) with stylized time course shown at the bottom displaying both transient and sustained elements. We allow for a variable degree of input coherence across V1 and LM (‘input consensus’ dial). **(B-D)** Dynamics of V1-LM consensus for stimulus detection. **(B)** Network activity projected onto the ‘agree’ (green) and ‘disagree’ (purple) modes of balanced, unselective activity (green and purple insets; recall Figure 4B), in response to a ⊘ stimulus that drives V1 while suppressing LM albeit less strongly (gray inset). Response projections are shown for three values of the specific long-range connection parameter 𝓁_s_ (0, 0.5 and 0.9), with diamond marks indicating the point of maximum consensus and triangular marks indicating 200ms after that. **(C)** Same data as in (B), with the projection of momentary, trial-averaged network responses onto the ‘agree’ mode (green line in B) now plotted against its ‘disagree’ counterpart (purple line in B). Diamond and triangular marks as in (B). **(D)** Same data as in (B-C), now showing the projections onto *local* unselective modes (presence vs. absence of stimulus) in V1 and LM against each other. **(E-G)** Same as (B-D), for stimulus discrimination. In this case, V1 is strongly driven by a ⊘ stimulus whilst LM is more weakly driven by a ⊘ stimulus (gray inset). The relevant agree/disagree modes are now the selective modes (green and purple insets), corresponding to consensus about the identity of the stimulus, rather than its presence/absence. As for detection, this conflicting stimulus gives rise to the correct consensus (⊘) especially for large 𝓁_s_. This happens even though the input itself presents more disagreement than agreement (F, black).

Likewise, when the input to both areas has the same total magnitude but is conflicted about stimulus orientation (Figure 5E, gray inset), specific long-range connections contribute to the emergence of a consensus about stimulus identity (Figure 5E-G). Importantly, this consensus favors the alternative that is more strongly supported by the input (here, +45°; see Figure 5G, dashed, for the opposite scenario). This is true even when the stimulus contributes more to the disagree-selective than to the agree-selective mode (as in the case shown here).

## Discussion

Here, we set out to elucidate the role of long-range connectivity in orchestrating dynamics across cortical modules. By combining data-driven and mechanistic modelling, we developed latent circuit models of observed neural activity across mouse V1 and LM, which were constrained by known properties of cortical circuit organization. These models uncovered slow reverberation of activity through long-range connections between the two areas. Further mathematical modelling re-vealed how this dynamical motif constrains the activity of distributed cortical modules in a way that ensures consistency of computation, or ‘consensus’ between them.

### Issues with model identifiability and how to mitigate them

Identifying dynamical interactions between brain areas from concurrent observations of their activity is in general an ill-posed problem. Indeed, when trying to account for observed neural activity using a network model, it is difficult to unequivocally tease apart external and recurrent contributions to the input that drives each neuron’s fluctuations [Pandarinath et al., 2018, Schimel et al., 2022, Malonis et al., 2021, Soldado-Magraner et al., 2023], as neither input is directly observed. In principle, even when using rich single-trial data, no approach is immune to wrongly inferring a mechanism not actually present in the cortical circuit [Qian et al., 2024, Genkin and Engel, 2020]. Our approach mitigates this concern in two ways. First, we include responses to optogenetic perturbations in the dataset used to fit the model; thus, the time course of at least some of the external inputs to specific cells is known in at least some of the trials. Indeed, such perturbations apply instantaneous, direct input to known cells, in contrast to e.g. sensory stimuli which enter the circuit of interest after largely unknown spatial and temporal filtering. Second, by introducing biological constraints into the model, we not only restrict the space of possible models that fit the recorded neural activity, but also expose the model to a series of experimentally testable validation criteria. For example, we were able to exclude models that would wrongly label known PV cells as excitatory, and explicitly simulate the effect of cell type-specific photo-activation to predict the corresponding neural responses. Finally, our model ultimately made qualitative predictions about the relative timescales of activity in different cross-area modes, which we were able to verify completely independently of our specific model fits (Figure 4).

### Generalizing to other mechanistic models

Mechanistic models of cortical circuits have classically focused on capturing the average behaviour of large neuronal populations, and have proven remarkably effective at explaining non-trivial qualitative features such as oscillations, global E/I balance, normalization effects, surround suppression, etc [Rubin et al., 2015, Kraynyukova and Tchumatchenko, 2018]. However, it remains unclear how these models should be extended to account for more detailed aspects of a circuit’s behaviour, and how their parameters could be constrained quantitatively using large-scale time series of neural data. Our work outlines a systematic path for distilling detailed recordings of large neuronal populations into the parameters of rich mechanistic models.

### Role of long-range connections in sustaining activity in the cortex

Our models and analyses make experimentally testable predictions. Specifically, we predict that stimulus-specific external input to the visual cortex is predominantly restricted to stimulus onset and offset, while the sustained cortical responses are supported by long-range cortical connections. Notably, the transient time course of our inferred external input resembles recent recordings from the visual thalamus (dLGN, Siegle et al., 2021). Paradoxically, despite the transient nature of feedforward thalamic input, intact thalamic activity was shown to be essential for sustained cortical responses: silencing the thalamus via optogenetic activation of the thalamic reticular nucleus (TRN) leads to a rapid decay of activity in V1 [Reinhold et al., 2015]. At first glance, this appears to also contradict our predictions. However, it is important to consider that TRN activation inhibits not only dLGN but also higher-order thalamic areas (e.g., pulvinar), which are thought to modulate corticocortical interactions [Sherman and Guillery, 2011, Saalmann and Kastner, 2011]. This could effectively isolate V1 from other cortical areas. Indeed, the rapid decay of cortical activity observed in Reinhold et al. [2015] is consistent with the fast decay time constants we identified in the isolated dynamics of our model’s V1 population. More broadly, beyond visual networks, sustained cortical activity in decision making or motor planning has also been shown to rely on multi-area interactions [Li et al., 2016, Guo et al., 2017].

### Role of long-range connections in consensus building

Here, we have found that the coupled dynamics of V1 and LM implement a form of consensus algorithm, whereby the two areas progressively get to reconcile their views about the presence of a stimulus and its coarse orientation. The fairly simple nature of this consensus arguably reflects the simplicity of our experimental go/no-go task. However, we hypothesize that dynamic consensus is a general feature of cortical dynamics that could play out at finer scales and be modulated to meet complex behavioural demands. Importantly, achieving fine-grained consensus would require detailed specificity in long-range connections between cortical areas. Just how such specificity could be achieved and regulated by behavioral context or learning is largely unknown. One possible mechanism would exploit trans-thalamic pathways, which appear to systematically mirror direct cortico-cortical pathways [Halassa and Sherman, 2019, Shepherd and Yamawaki, 2021]. Detailed gain modulation of thalamic neurons involved in those pathways (e.g. pulvinar, known to send functionally specific projections to V1; Furutachi et al., 2024) could provide sufficient flexiblity for regulating multiple modes of consensus between cortical areas. Indeed, Mo et al. [2024] showed that inhibiting the trans-thalamic pathway between primary and higher-order somatosensory cortices in mice leads to a loss of learning-induced texture selectivity, but no change in overall cell responsiveness to tactile stimuli. Our model of Figure 5 would attribute such effects to a decrease in *specific* longrange connectivity affecting consensus in the *selective* mode useful for stimulus discrimination, but not affecting the unselective mode useful for stimulus detection. More generally, richer forms of consensus arising from fine-grained connectivity could serve more complex computations, for example the integration and reconciliation of bottom-up sensory information with top-down prior expectations [Knill and Pouget, 2004]. By integrating data from large-scale functional connectomics [MICrONS Consortium et al., 2021] with multi-area neural recordings during more complex tasks, our theoretical approach is ideally positioned to test such hypotheses and uncover the richer dynamics of brain-wide consensus.

## Methods

### Experimental procedures

No new experimental data were collected for the purposes of this study. The acquisition and pre-processing of data used in this study are described in detail in Javadzadeh and Hofer [2022]. From the total of 14 mice included in Javadzadeh and Hofer [2022], we sub-selected 7 mice for inclusion in this study, based on the criterion that the electrophysiological recordings contained at least one well-isolated single unit that was identified by the optogenetic perturbations as PV+. Models were fit using all trial types, but only trials in which the mice performed the task correctly were included in subsequent analyses, unless specified otherwise. The spiking activity of the recorded neurons was binned at 5ms resolution, and for visualization, smoothed with a running average of 25ms or 5 bins (Figures 1B,C,G,2A,D,3C).

### Latent circuit model of V1/LM data

#### Latent circuit dynamics

We modelled latent circuit dynamics as an input-driven recurrent neural network described by a standard firing rate equation [Dayan and Abbott, 2005]. Specifically, the circuit’s *n*-dimensional ‘latent state’ ***z*** evolved according to

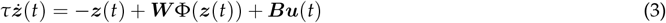

where *τ* = 20ms is a single-neuron characteristic time constant, ***W*** is a matrix of recurrent connectivity (see below), ***B*** is a matrix of input weights, and 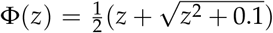 is a soft rectified-linear activation function. Note the presence of external inputs ***u***(*t*) described in detail below. The spiking activities of our *N* recorded neurons were then modelled as conditionally independent Poisson processes given the latent circuit’s activity, ***z***(*t*), with momentary firing rates ***r***(*t*) given by:

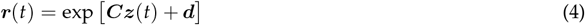

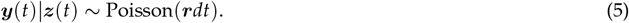

Here, ***C*** is a *N*× *n* matrix of output weights and ***d*** is an *N*-dimensional vector of constant offsets. Equation 3 was discretized using a time step *dt* = 5*ms*. All model parameters were optimized to fit the electrophysiological data (see below, ‘Network training procedure’). Critically, ***W***, ***C*** and ***B*** were constrained to reflect biophysical properties of the V1-LM network (see below; schematics in Figure 1D-F).

Note that Equation 3 does not include a constant input term. We found that including such a bias term caused the model to fall into local minima, consistently learning solutions with worse residual log-likelihoods (see Figure S1E).

#### External inputs

Our model captures trial-by-trial variability in neural activity not only via the Poisson sampling step in Equation 5, but also – and more importantly – through trial-by-trial fluctuations in the external inputs ***u***(*t*). These (deterministically) produce variations in latent circuit activity according to Equation 3, and therefore also in the neurons’ firing rates (Equation 4). In the language of probabilistic modelling, the external inputs ***u*** constitute the model’s latent variables.

Simultaneously inferring dynamics *and* external input is a fundamentally ill-posed problem, which our probabilistic model addresses by placing task-informed, non-stationary prior distributions on the latent inputs. Specifically, we used three input channels – i.e. ***u***(*t*) ≡ [*u*_0_(*t*), *u*_1_(*t*), *u*_2_(*t*)]^⊤^, each entering the latent circuit through input weights given by the corresponding column of the *n* × 3 matrix ***B*** (Equation 3). For each input channel *i*, we assumed *u*_*i*_(*t*) to be (a priori) *independently* and normally distributed across time steps – ensuring that any continuous/smooth fluctuations in firing rates could only be accounted for by recurrent dynamics in the latent circuit. Moreover, the variance of this Gaussian prior was given a channel- and trial-specific temporal profile reflecting the known timing of the corresponding stimulus:

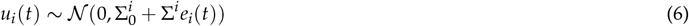

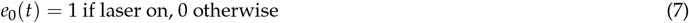

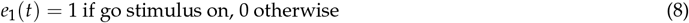

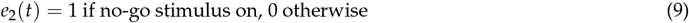

where 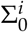 and 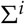 are two positive variance parameters optimized alongside all other model parameters (see below).

Given that the laser input in our experiments had a direct effect only on inhibitory neurons, we constrained the first column of ***B*** (associated with *u*_0_(*t*)) to be zero for all sub-populations except for the inhibitory neurons of the targeted area. Additionally, we ensured that the weights of this column of ***B*** were all positive. Finally, to eliminate the degeneracy that exists between the scale of the inputs ***u***(*t*) (set by 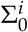 and 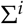 as detailed above) and the scale of the matrix ***B***, we constrained the norm of each column of ***B*** to be equal to 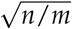 (where *n* is the number of units in the latent circuit, and *m* = 3 is the number of input channels). The ***B*** matrices learned for all animals are shown in Figure S2.

#### Constraints on the latent circuit connectivity

We partitioned the latent circuit’s activity ***z***(*t*) into two halves, corresponding to the V1 and LM subcircuits respectively (see Figure 1D). Within each subcircuit, we took the first half of the latent units to be excitatory, and the other half to be inhibitory. This partitioning of the circuit into four sub-populations allowed us to enforce Dale’s law, as well as the purely excitatory nature of long-range projections, by constraining the recurrent weight matrix ***W*** to have the following structure:

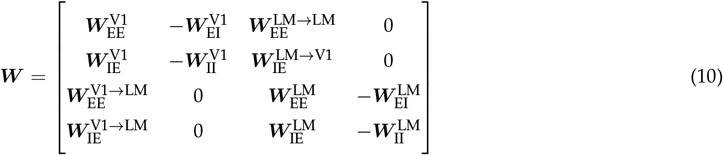

with all elements of the various 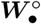 blocks constrained to be positive. We enforced the sign constraints in our model by passing elements of ***W*** through a positive nonlinearity, and multiplying ***W*** with a mask matrix containing the sign of each element. We note that, in related work, Jha et al. [2024] proposed a method to learn linear latent dynamical systems constrained to follow Dale’s law using a constrained quadratic optimization approach.

#### Structured sparsity constraint on the latents-to-neurons readout

The matrix ***C*** in Equation 4, which determines how the firing rates of the recorded neurons (corresponding to rows) are assembled from the activity of the latent units (corresponding to columns), was constrained such that the neurons recorded in V1 (resp. LM) would only be associated with V1 (resp. LM) latent units. This was achieved by enforcing the following block structure (see Figure 1F):

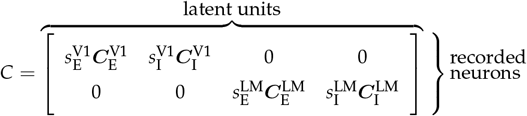

where each 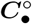 is an element-wise positive matrix with unit-norm columns, and each corresponding 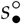 is a positive scalar. This per-block column-wise normalization of ***C*** balances the model internally by ensuring that all the latent units within each sub-population have a comparable effect on the activity of the observed neurons. Moreover, the inclusion of separate scale factors 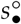 allows the different E/I sub-populations to contribute to different degrees to the neural activity.

Importantly, to facilitate interpretability of the latent circuit, we learned the model in such a way that it would unequivocally label each recorded neuron as being excitatory or inhibitory. We achieved this by included in the overall cost function (see below) a structured sparsity penalty on ***C*** that encourages each recorded neuron to be locally associated either with the excitatory latent units, or with the inhibitory latent units, but not with both types simultaneously. In other words, this penalty promotes parameter solutions in which the rows of ***C*** are non-zero either within the 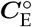 block or within the 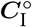 block (where ∘ denotes the relevant cortical area). This penalty took the following form:

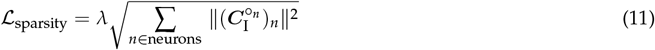

where °_*n*_ ∈ {V1, LM} is the cortical area where neuron *n* was recorded, and 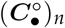 denotes the *n*^th^ row of the matrix block 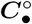, and ∥ · ∥ denotes the *L*_2_ norm. The scalar *λ* was set to 10^3^ following a hyperparameter search.

#### Definition of putative excitatory and inhibitory cells

For models trained with the above constraints, we were able to assign each neuron a unique excitatory or inhibitory identity based on the learned readout matrix, ***C*** (see Figure 1D). For each neuron, we calculated the *L*_2_ norms of the corresponding readout weights originating from the excitatory and inhibitory latent sub-populations *separately*, and labelled the neuron as E or I according to which of the two norms was the largest.

### Network training procedure

Our latent circuit model, together with the prior distribution over external inputs and the Poisson observation noise model described above (Equations 3, 5 and 6), constitute a probabilistic generative model whose parameters we directly optimized to fit our spiking data. To this end, we used iLQR-VAE [Schimel et al., 2022], a generic control-based algorithm for learning probabilistic, input-driven latent dynamics from neural population recordings. iLQR-VAE learns model parameters *θ* that maximize a lower bound on the log likelihood of the data, log *p*_*θ*_ (***y***). This evidence lower bound (ELBO; Kingma and Welling, 2013) is a standard objective, used when the true log likelihood cannot be evaluated in closed-form, as is the case in our model. The ELBO, denoted by L, relies on an approximate posterior distribution over inputs, *q*_*ϕ*_(***u***|***y***):

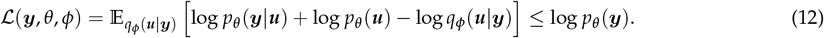

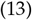

In iLQR-VAE, *q*_*ϕ*_(***u*** |***y***) = 𝒩 (*µ*(***y***), Σ) is parametrized as a Gaussian distribution, whose mean *µ*_*θ*_ (***y***) is defined as the most likely set of inputs given the data and the model parameters. This maximum a posteriori estimate can be efficiently obtained using the iLQR algorithm [Li and Todorov, 2004]:

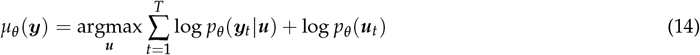

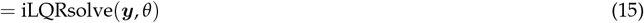

As in Schimel et al. [2022], we defined the covariance **Σ** as a trial-independent, separable matrix, i.e as the Kronecker product of a spatial factor **Σ**_s_ and a temporal factor **Σ**_t_, which were learned throughout training and shared across all training trials.

In summary, fitting our latent circuit model to the V1-LM spiking data involved jointly optimizing all model parameters *θ* and the approximate posterior parameters *ϕ* = {*θ*, **Σ**_s_, **Σ**_t_} to minimize the following combined objective:

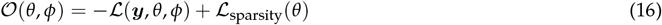

### Log-likelihood computations

#### Computation of cross-validated log-likelihoods

To validate the performance of our model, we computed its ability to predict the activity of held-out neurons, given firing rates inferred using the held-in neurons. We held out one neuron at a time. To predict the activity of held-out neuron *j*, we inferred inputs as 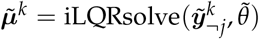, where 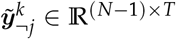 is the spike trains of all neurons, excluding neuron *j*, in trial *k* (and 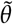 are the model parameters with the *j*-th row of ***C*** and ***d*** masked out). We then computed the predicted firing rates for all (both held-in and held-out) neurons 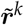 by unrolling the trajectories induced by the inputs 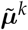 (using the full set of parameters *θ*). In turn, this allowed to compute the log-likelihood of the spikes in trial *k* for the held-out neuron *j*, as

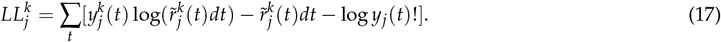

#### Computation of the empirical log-likelihood

As a baseline to compare the model predictions to, we computed the empirical log-likelihood for a trial *k* by evaluating the predicted activity for every neuron using that neuron’s average activity across all the other trials from the same condition *c*, leading to a predicted firing rate time course

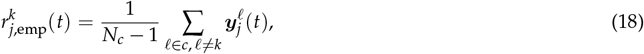

where *N*_*c*_ is the number of trials in condition *c*. Given these empirical firing rates, we computed the empirical log-likelihood for neuron *j* at trial *k* as

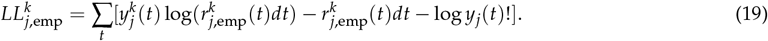

#### Residual log-likelihood

We define the residual log-likelihood for a given neuron *j* as 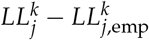. If this quantity is positive, it means that the prediction of the model for that neuron is more accurate than a prediction based on trial averaging, i.e., that the model is able to capture meaningful single-trial variability in the data. Residual likelihoods were calculated separately for each neuron across 18 different conditions (2 visual stimuli and 9 silencing condition for each visual stimulus), and then averaged across all trials and conditions.

### Model selection

#### Choice of hyperparameters

To select the model hyperparameters *n* and *m* (number of latent state variables and input channels, respectively), we used a 3-fold cross-validation approach. For each animal, we split the trials into 3 subsets. Then, for each possible pair of subsets among these three, we trained a model using the data from that pair and subsequently computed the heldout log-likelihoods on the remaining subset. Finally, we averaged the results over the three pairs, over animals, and over neurons.

We first selected the optimal value of *n* for the model with three input channels (*m* = 3) corresponding to the visual and optogenetic stimuli, as described above. We explored model sizes ranging from *n* = 8 to *n* = 24 in increments of 4, and selected the minimal value of *n* after which the residual log-likelihood stopped improving (see Figure S1A). Having selected and fixed the optimal value for *n*, we checked whether the choice *m* = 3 was optimal, using the same model selection procedure. When varying the number of input channels, we considered both (i) having multiple channels corresponding to each prior variance profile (c.f. Equations 7 to 9), i.e. multiple channels for each external stimulus (Figure S1B), and (ii) the addition of channels with temporally unmodulated prior variance (see Figure S1C). Neither of those increased model performance relative to using *m* = 3, which is the minimal number of channels allowing to have one input per external stimulus. Note that models with additional input channels could in theory capture timing difference in the visual input to V1 and to LM. However, we found that having one channel per input yielded the best performance on the validation fold.

Additionally, we compared our models to models with the same architecture but for which the inputs were not inferred, and were instead fixed to follow the envelope corresponding to each external stimulus. This implied that their time course was constrained to be the same for every trial of a given condition. Those models performed considerably worse than the models with inferred inputs (Figure S1D). Other model hyperparameters such as the spectral radius of ***W*** at initialization and the Adam learning rate were fixed to values that allowed robust training. Final hyperparameter choices are reported in Table 1. All trials, irrespective of behavioral outcome, were included for log-likelihood calculation and model selection.

**Table 1:** Model hyperparameters.

| symbol | value | unit | description |
| --- | --- | --- | --- |
| $n$ | 16 | - | number of latent units |
| $m$ | 3 | - | number of latent inputs |
| $\tau_E$ | 20 | ms | excitatory latents time constant |
| $\tau_I$ | 20 | ms | inhibitory latents time constant |
| $\eta$ | 0.004 | - | learning rate |
| $k$ | 10 | - | scaling of the optimizer square root decay |
| $r$ | 0.6 | - | spectral radius of $\mathbf{W}$ at initialization |
| $\lambda$ | 1000 | - | scale of the regularization for $\mathbf{C}$ |
| $F_e$ | 0.5 | - | fraction of excitatory neurons |

#### Selection of models for plotting and analysis

For the constrained models, having set the hyperparameters as described above, we trained 10 models with different random seeds (i.e. different random initializations of the model) per animal. This was done to reduce the chance of getting stuck in local minima. Moreover, as our conclusions were dependent on the learned values of the long-range weights, and to avoid biasing our models, we varied the value of the long-range weights at initialization. More precisely, we varied the ratio of the norm of long-range weights to local weights at initialization between 1 and 1.6 in steps of 0.2. We discarded models that diverged during training (41 out of 280 models in total). Out of the remaining models, we then picked the best model for each animal, across initialization seeds and long-range weights, for further analyses and plotting. For each animal, the best model was selected by first sub-selecting the models that classified the known PV cells correctly as inhibitory (187 out of 239, i.e. 78.24% of the models; see Figure S1F). Among these, we picked the model that yielded the highest cross-validated log-likelihood. Furthermore, we only included active cells (neurons whose spike count during the stimulus in control trials had a signal-to-noise ratio, i.e. mean/std over trials, larger than 1) for log-likelihood calculations.

For the unconstrained models, we used 5 random initialization seeds. However, as inhibitory cell identities were not defined in these models, we picked the best model based only on the held-out log-likelihood criterion explained above.

### Calculating covariances

In Figure S3, we calculated *N* × *N* noise covariance matrices in both data and model-predicted activity as:

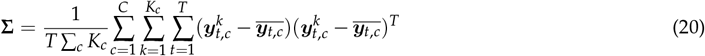

where *c* indexes conditions (2 visual stimuli and 9 silencing condition per stimulus), 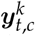 is a *N* × 1 vector denoting spike count of *N* neurons in 25ms bins, in control condition *c* (no optogenetic stimulation), trial *k*, and time *t* (*K*_*c*_: number of trials in condition c,*T*: number of time points, *N*: number of neurons). 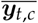 is the trial-average activity in condition *c*. For calculating model covariances, we sampled pseudo-observations ***y*** from a Poisson distribution whose mean was taken to be the posterior predicted firing rates. All trials, irrespective of behavioral outcome, were used for calculating covariances. Variances in Figure S3 are the diagonal values of **Σ** and cross-covariances are its off-diagonal values.

### Linearization of the dynamics

Around a (approximate) fixed point ***z*** _*f*_, the dynamics in Equation 3 can be Taylor-expanded to first order, leading to a linear dynamical system whose dynamics matrix is given by the Jacobian ***A***:

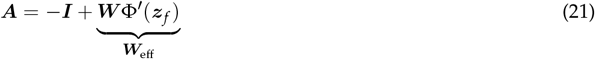

Here, ***W***_eff_ can be thought of as a matrix of “effective connectivity”.

For a given trial *k*, we defined 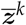 as the time-averaged activity either before or during stimulus, i.e 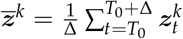 with Δ = 400ms, *T*_0_ = − 400ms for the pre-stimulus window and *T*_0_ = 100ms for the stimulus window (*T*_0_ is measured relative to visual stimulus onset). This choice was motivated by the fact that the dynamics exhibited very small velocities in these time windows; see Figure 3E. We then defined 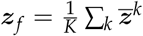.

### Computation of the dynamics distance

In Figure 2C, we computed the similarity between the dynamics of the model for different animals, as a normalized Procrustes distance (see Williams et al., 2021 and Ostrow et al., 2023) between their linearized dynamics, i.e as :

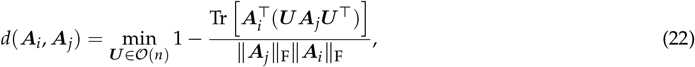

where ***A***_*i*_ and ***A***_*j*_ denote the linearized learned dynamics for animals *i* and *j* (obtained as described in Methods - Linearization of the dynamics), ∥ · ∥_F_ denotes the Frobenius norm, and ***U*** is an orthogonal (rotation) matrix (optimization over ***U*** is necessary in order to account for the fact that the dynamics may be equivalent up to a rotation). We used the average distance *d*(***A***_*i*_, ***A***_*j*_) across all pairs of animals (i.e. all (*i, j*) such that *i > j*), as our measure of consistency of the learned dynamics across animals.

As shown in Ostrow et al. [2023], *d*(·,·) is a valid distance metric, bounded between 0 and 1, which computes the similarity of the vector fields of two dynamical systems. While Ostrow et al. [2023] applied this analysis to dynamical systems identified via delay embedding of the dynamics, we instead apply it directly to the linearized dynamics of our model.

To perform the minimization in Equation 22, we parametrized the orthogonal matrix ***U*** using a Cayley transformation [Ostrow et al., 2023]. As pointed out in Ostrow et al. [2023], the optimization landscape is disjoint for ***U*** matrices with det ***U*** = 1 and det ***U*** = −1. Thus, for each pair of dynamics matrices, we perform the optimization over matrices ***U*** such that det ***U*** = 1 as well as over matrices ***U*** such that det ***U*** = − 1, and use the minimum distance across those two subsets.

### Comparing the model’s ability to capture the effect of optogenetic perturbations

To evaluate how well the models captured the effect of simulated optogenetic perturbations (Figure 2D), we first evaluated the inferred input for each no-go, no-laser trial. We then ran the model dynamics forward with those inputs, whilst additionally perturbing the inhibitory model population in either V1 and LM, depending on which area expressed ChR2 in our experiments (different across animals). Note that these perturbations were simulated for each trial, based on the inferred input for that trial. We could then compare the neural responses predicted by the latent circuit model to the corresponding photo-stimulation responses observed in the experiments. Specifically, we used a simulated pulse of optogenetic input modeled as

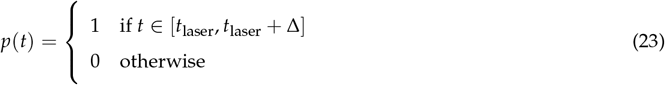

where *t*_laser_ is the onset time of laser stimulation in the relevant silencing condition in the experiments, and Δ = 150 ms is the laser duration. We assumed that this input influenced the latent units via a weight vector *B*_*p*_, whose elements were non-zero only for the inhibitory latent units of the stimulated area. We optimized the non-zero elements of *B*_*p*_ to maximize the log-likelihood for the spike trains of the known PV cells in the relevant perturbation trials. We then measured the average predicted perturbation-induced change (relative to no-perturbation), 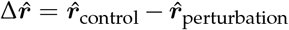, in the rest of the neurons during the stimulation time window, and compared it to the same quantity, Δ***r***, measured in the data. We report the quality of fit as the Pearson correlation between 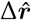 and Δ***r***. This is plotted for one animal in Figure 2D middle, and for the rest of the animals in Figure S4.

As a comparison (see Figure 2D), we repeated the above for “control-only” models which were trained on control trials without optogenetic perturbation (2/3 of the control trials were used for training). We trained a minimum of 12 models per animal, and chose the best model following the same procedure used for the default models; see Selection of models for plotting and analysis. (For one of the animals, no model —out of 30 models— resulted in correct classification of all PV neurons; for that animal, we only used the log-likelihood criterion for model selection.)

### Spike width histograms

We extracted the average spike waveforms for each neuron, and the spike width was defined as the width of this waveform at 10% of its full amplitude (Figure 2E).

### Analysis of the role of inhibition in the dynamics

To evaluate the role of inhibition in stabilizing the dynamics (Figure 2F), we measured the stability of our latent circuit dynamics, in the presence or absence of inhibition. We measured stability before and during stimulus presentation by computing the effective connectivity ***W***_eff_ (see Equation 21 in Methods - Linearization of the dynamics). We then computed the largest real part of the eigenvalues of the effective linear dynamical system, *λ*_*max*_ = max_*i*_(ℜ(*λ*_*i*_)) where *λ*_*i*_ are the eigenvalues of ***W***_eff_. A (linearized) network is said to be “inhibition-stabilized” if *λ*_max_ *<* 1 (stable) when computed on the full ***W***_eff_, but *λ*_max_ *>* 1 (unstable) when all the inhibitory weights in ***W***_eff_ are set to zero.

### Connectivity strength as a function of the noise correlation

In Figure 2G, we computed the noise correlation matrix of the mean-subtracted latent circuit responses of the V1 excitatory subcircuit during control no-go trials, ***z***_*t*_, as follows :

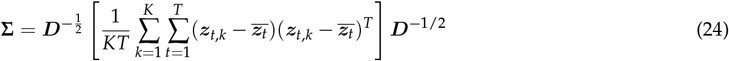

where *t* and *k* denote time bin and trial, respectively, 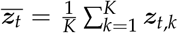 and ***D*** is a diagonal matrix of single-neuron variances, i.e. 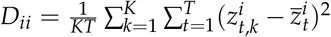.

In Figure 2G, we plot the effective connection weights, 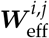 (computed based on the stimulus period as described by Equation 21), for pairs of excitatory latent units (*i < j*), as a function of the pairs’ noise correlation **Σ**^*i,j*^ (more specifically, we binned pairs based on their noise correlation and created box-whisker plots of the effective connection weights for each bin). When comparing with the circuit at initialization (Figure 2G, top), we considered ***W***_eff_ and ***z***_*t*_ to be the connectivity matrix and latent trajectories at the first iteration of training the model. We repeated the same procedure for go trials, with similar results (Figure S5).

### Calculating recurrent and external currents

For analyses described in Figure 3A-B, we defined external and recurrent currents as 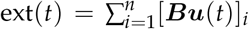 and 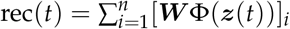, respectively.

### Sensitivity of the networks

In Figure 3A, B, and J, we evaluated the sensitivity of the latent circuit to changes in the inputs vs. changes in the recurrent weights by running the network dynamics forward, using the inputs inferred from the data for every test trial, but including a gain *γ* that we used to either scale down the input matrix ***B*** (see Equation 25), or the connectivity matrix ***W*** (see Equation 27):

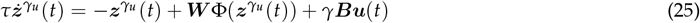

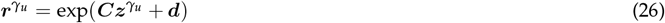

vs.

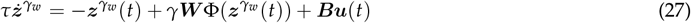

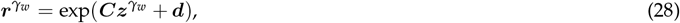

We computed the sensitivity by measuring changes in the total activity in no-go trials, either before or during stimulus onset, and normalizing those to the activity obtained for *γ* = 1, i.e. 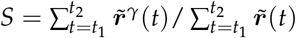 where 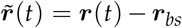, with ***r***_*bs*_ the average baseline (pre-stimulus) activity.

We used the same approach to compute the sensitivity separately to either local or long-range weights, which was done by applying the gain to the corresponding local (*W*^*LM*^ and *W*^*V*1^) or long-range blocks (*W*^*LM*→*V*1^ and *W*^*V*1→*LM*^) of the ***W*** matrix (see Equation 10).

### Intrinsic flow and velocity

In Figure 3D, we plot the velocity field of the intrinsic dynamics (i.e dynamics in the absence of external inputs), projected onto the subspace spanned by the top two principal components (PCs) of the latent trajectories. Note that projections onto the PCs were only used for visualization purposes, and all analyses were performed using the full-dimensional dynamics. We first performed a singular value decomposition on the trial-averaged latent activity in no-go trials *Z* ∈ R^*n*×*T*^ as *Z* = *U*Σ*V*_*T*_, before defining 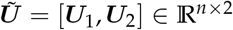 as the top 2 PCs. We then computed the projected velocity field at each point in the 2D space, ***x*** = (*x, y*), as ***v***(***x***) ∈ R^2^, where:

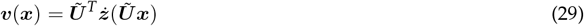

and the function ***ż*** (·) is given by:

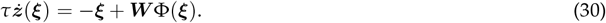

To compute the velocity in Figure 3E, we similarly used Equation 30, but we used the no-go trial-averaged latent trajectories (without projecting onto PCs) for ***ξ***. In Figure 3H, we followed the same procedure, but using the ***Z*** and ***W*** restricted to each area. In this case, the 2 PCs were similarly extracted from the area-restricted latents.

### Network time constants and line attractor score

For analyses in Figure 3F, G, I, and J, we linearized the dynamics around the average value of the latents across trials and time, either during or before the stimulus, and computed the eigenvalues and eigenmodes of the linearized dynamics ***A*** (see Methods - Linearization of the dynamics). In this continuous-time linear dynamical system, each eigenmode *j* evolves in time according to 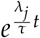, where *τ* is the single neuron time constant and *λ*_*j*_ the eigenvalue of mode *j*. The characteristic decay timescale of each mode is given by 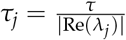. Assuming the modes are ordered such that 0 *>* Re(*λ*_0_) … ≥ Re(*λ*_*n*_), i.e *τ*_0_ ≥ … ≥ *τ*_*n*_, *τ*_0_ defines the slowest timescale in the dynamics.

To calculate the time constants in the V1-only or LM-only networks in Figure 3I, we followed the same procedure but used ***W*** and ***Z*** restricted to each individual area to compute the linearized dynamics. When comparing the time constants of these single-area networks to the full network, in order to control for their smaller size, we constructed subnetworks of the size of each individual area (i.e. of size *n*/2), sampled randomly from the full latent network (500 random subsets, and excluding any subselection that would correspond to the V1 or LM network).

To quantify the existence of a line attractor in the dynamics, we compute the “line attractor score”, defined as in Nair et al. [2023] as a log ratio of the slowest to the second-slowest time constant of the network dynamics, i.e. log(*τ*_0_/*τ*_1_)/ log 2. A true line attractor would correspond to an infinite line attractor score. A score of 1 means that the slowest mode is twice as slow as the next mode. A score of 0 means that the first two slowest modes have the same time constant (as happens, e.g., when these two modes define a plane with rotational dynamics, i.e, the imaginary parts of their eigenvalues are non-zero).

### Minimal E-I networks

Our minimal E/I networks (Figures 4 and 5) are described as linear rate models, consisting of two areas, where each area’s connectivity is given by:

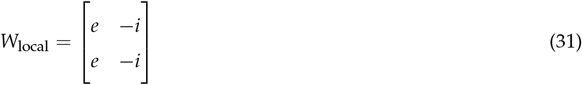

where *e* and *i* are the strength of excitatory and inhibitory connections, respectively. The activity in the full network evolves as:

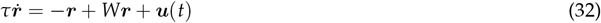

where ***u***(*t*) is an external input which is zero unless otherwise specified,

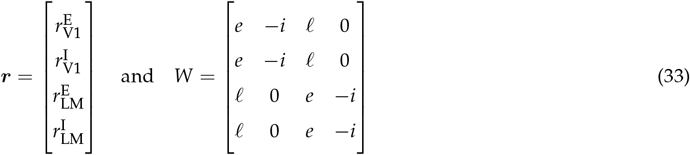

and 𝓁 is the strength of long-range excitatory connections. This is the minimal network architecture depicted in Figure 4B.

We can show that the orthonormal basis *Q* consisting of vectors *Q* = [***b***_***a***_, ***u***_***a***_, ***b***_***d***_, ***u***_***d***_] (‘*b*’ for ‘balanced’, ‘*u*’ for ‘unbalanced’; ‘*a*’ for ‘agree’, ‘*d*’ for ‘disagree’), where:

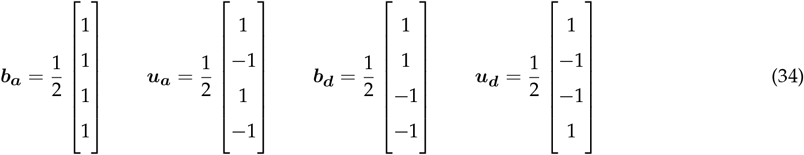

is a Schur basis of *W* such that 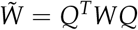 is the upper triangular matrix

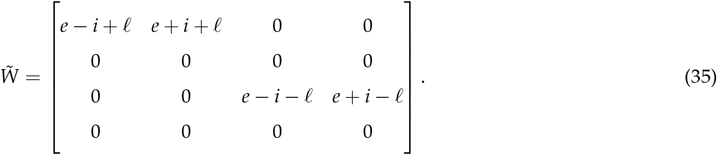

This upper triangular form describes feedforward connectivity in the new basis *Q*, and reveals the existence of two separate functional subnetworks, respectively describing the dynamics of agreement and disagreement between V1 and LM. The dynamics of each functional subnetwork are characterized by balanced amplification due to the feedforward weight from the unbalanced to the balanced mode [Murphy and Miller, 2009]. See Supplementary Section S2.1 for more details, including the interpretation of different elements of the Schur form.

We also considered a version of the above minimal model that also incorporated a notion of selectivity for go vs. no-go stimuli (Figure 4C). Specifically, we split every E and I population in V1 and LM into two sub-populations, one receiving direct go input, and the other receiving direct no-go input. This resulted in a circuit with 8 units, with recurrent connectivity parameterized as:

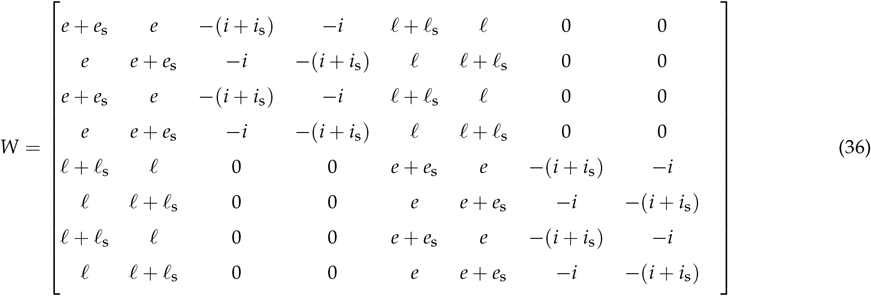

where *e*_*s*_, *i*_*s*_, and *l*_*s*_ control the stimulus selectivity of local excitatory and inhibitory, and long-range excitatory connections, respectively. The Schur decomposition of this network, along with its interpretation in terms of time constants, can be found in Section S2.

### Autocorrelation of neural data

The autocorrelation of the agree (*a*) and disagree (*d*) neural activity patterns across V1 and LM are defined, respectively, as the sum and difference of the average empirical spike counts binned at 5 ms, *s*(*t*), within each area, i.e.,

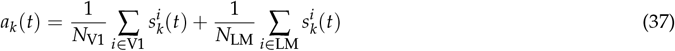

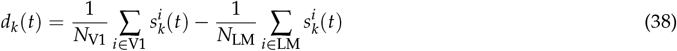

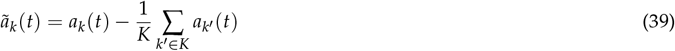

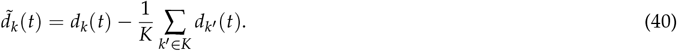

We define the autocorrelation of the agree mode as the autocovariance normalized to the overall variance:

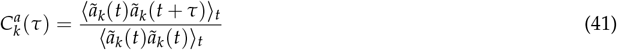

where ⟨·⟩_*t*_ denotes an average over time bins *t* that are such that both *t* and *t* + *τ* fall within the relevant time window. This time window was [−400, 0] ms (‘pre’) or [100, 500] ms (‘during’) relative to stimulus onset. The autocorrelation of the disagree mode, 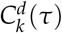, is defined analogously. See Figure S7B for a distribution of marginal variances (denominator in Equation 41) in the agree and disagree modes.

In Figure 4H and I, we report the mean autocorrelation and its standard error across all correct control go and no-go trials and all animals. Note that mean subtraction in Equation 39 was done separately per animal/condition for go and no-go trials.

Our minimal models of V1-LM dynamics (Figure 4B-C) also make predictions for the decay timescales of balanced vs. unbalanced modes. Estimating the autocorrelation time constant of these modes required estimating the E/I identity of each recorded neuron. For this, we used the identities inferred by the latent circuit models, and computed the momentary contributions of the balanced and unbalanced agree (*a*_*b*_, *a*_*u*_) or disagree (*d*_*b*_, *d*_*u*_) modes to the recorded activity as:

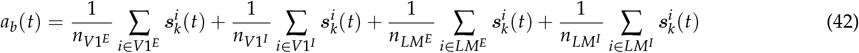

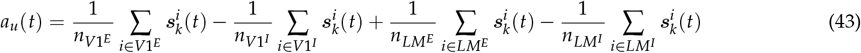

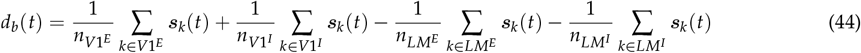

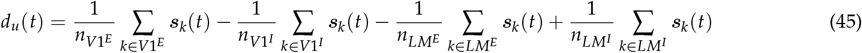

Having defined these projections, we follow the same procedure as in Equation 39 - Equation 41 to calculate autocorrelations. Results are shown in Figure S7A

### Quantifying autocorrelation differences

To quantify the difference between the autocorrelation functions of neural activity projected onto the agree vs. disagree modes (Figure 4H and I), we first identified the time lag at which the average agree/disagree autocorrelation function reached its maximum (*t*_max_). We then quantified the difference, denoted by Δ_max_, between the agree and disagree autocorrelation functions at this time point (see the caption of Figure 4).

### Projected autocorrelations for selective/unselective modes

In Figure 4I, to estimate the time course of the selective and unselective modes from the neural data, we computed two indices for each neuron. The first index measured ‘unselective’ responsiveness, i.e. how much more each neuron responded to either stimuli (go/no-go), relative to baseline. The second index measured ‘selective’ responsiveness, i.e. how much each neuron preferred the go stimulus over the no-go stimulus. This resulted in two vectors of indices:

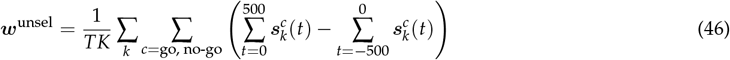

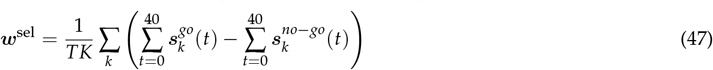

We then used these indices to compute weighted averages of the neural activity at each time step, 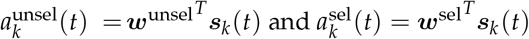.

The distributions of unselective and selective weights (***w***^unsel^ and ***w***^sel^) across V1 and LM neurons are shown in Figure S7D-E, and their relationship with one another is shown in Figure S7C. The elements of ***w***^unsel^ were biased towards positive values (Figure S7D), as most recorded neurons responded to visual stimuli by increasing their firing rates. For ***w***^sel^, in contrast, the distribution was symmetric. This is because we measured responses early during stimulus presentation, i.e. likely before any go-stimulus-related behavior could break the symmetry in the neural responses to the two stimuli (Figure S7E). The choice of a small time window to compute ***w***^sel^ also ensured that the selective index for a neuron was not corrupted by its unselective stimulus responsiveness, i.e that ***w***^sel^ were not directly correlated with ***w***^unsel^ (Figure S7C). This was important to establish that the slow time constant of the autocorrelation in agree-selective mode was not simply due to the correlation of this mode with the agree-unselective one, but instead depended on the stimulus selectivity of neurons.

### Alignment of the latent circuits’ eigenmodes onto agree unselective/selective modes

For our latent circuit models, we could ask whether their two agree modes (unselective and selective) bore any relationship with their two slowest eigenmodes. Eigenmodes were computed for the Jacobian of the latent circuit dynamics linearized around the average activity in no-go trials during stimulus presentation (see Methods - Linearization of the dynamics), and were sorted from slowest to fastest according to their associated eigenvalues. Similarly, we could estimate the latent circuit’s unselective and selective agree modes by computing, for each latent unit, similar indices of unselective/selective responsiveness as we had computed for recorded neurons in Figure 4I (c.f. Equations 46 and 47). For the selective indices, activity of the latent circuit during the onset period (0-100ms from the stimulus presentation) was used, as the latent circuit activity was strongly driven by external inputs during this time window (Figure 3A). This yielded two normalized vectors, whose overlaps with the eigenvectors we evaluated, by calculating the absolute value of their dot product. These overlaps are shown in Figure S7F.

### Details of Figure 5

In Figure 5, we simulated the linear dynamics of the minimal circuit model of Figure 4C, i.e. Equations 32 and 36 with parameters *τ* = 10 ms, *e* = 2, *i* = 2, *e*_s_ = 0, *i*_s_ = 0, 𝓁 = 0, and 𝓁_s_ taking values in the set {0, 0.5, 0.9}. The input to the network was ***u***(*t*) = *α*(*t*/*τ*^′^)***u***_0_ with *τ*^′^ = 15 ms, i.e. it was the product of a scalar temporal envelope (Figure 5A, bottom) 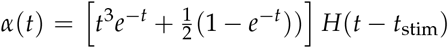 (where *H*(·) denotes the Heaviside function) and a spatial input pattern ***u***_0_ which expressed how much each subpopulation was driven by the ‘visual’ stimulus. For Figure 5B-D, we set ***u***_0_ = (1, 0, 1, 0, −0.6, 0, −0.6, 0)^*T*^, whereas for Figure 5E-G we set ***u***_0_ = (1, 0, 1, 0, 0, 0.6, 0, 0.6)^*T*^ (c.f. gray insets).

### Statistics

We used two-sided Wilcoxon rank-sum tests for independent group comparisons, and two-sided Wilcoxon signed-rank tests for paired tests, unless otherwise stated.

## Acknowledgments

This work was performed using resources provided by the Cambridge Service for Data Driven Discovery (CSD3) operated by the University of Cambridge Research Computing Service (www.csd3.cam.ac.uk), provided by Dell EMC and Intel using Tier-2 funding from the Engineering and Physical Sciences Research Council (capital grant EP/T022159/1), and DiRAC funding from the Science and Technology Facilities Council (www.dirac.ac.uk). MS was funded by an Engineering and Physical Sciences Research Council (EPSRC DTP) studentship (RG94782). SBH and MJ were supported by the Sainsbury Wellcome Centre core grant from the Gatsby Charitable Foundation and the Wellcome Foundation (090843/F/09/Z), and a Wellcome Investigator Award (S.B.H., 219561/Z/19/Z). MJ was additionally supported by the Cold Spring Harbor Laboratory Fellows Program. YA was supported by UKRI Biotechnology and Biological Sciences Research Council research grant BB/X013235/1. We would like to thank Ari Benjamin, Kyle Daruwalla, YoungJu Jo, and Ivan Voitov for feedback on the manuscript.

## Author contribution

MJ, MS, YA, GH conceived the study. MJ, MS, GH performed the analyses with feedback from YA and SBH. MJ, MS, GH, YA wrote the manuscript with feedback from SBH.

## Data and code availability

Data and code are available from the corresponding authors upon request.

