## Supplementary material for "Dynamic consensus-building between neocortical areas via long-range connections"

#### Supplementary figures

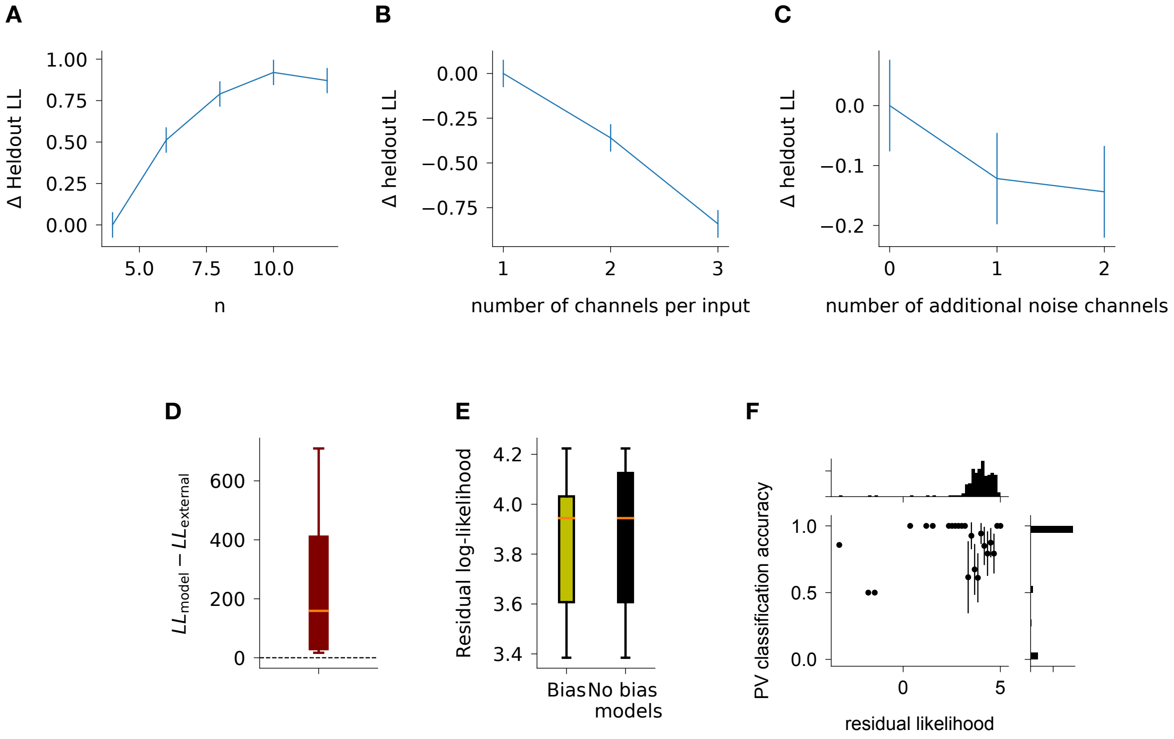


**Figure S1: Model selection** – **(A)** Normalized residual heldout log-likelihood of models of different size ($n$). Normalization was done by subtracting the average residual log-likelihood (across animals, cells, and cross validation folds) of the smallest model ($n = 4$). **(B)** Normalized residual heldout log-likelihood of models of size $n = 16$, as we increase the number of input channels. In this case, we used a stimulus-dependent prior, and used either $m = 1$, $m = 2$, or $m = 3$ input channels per stimulus (go, no-go, and opto). As in (A), normalization was done by subtracting the average residual log-likelihood (across animals, cells, and cross validation folds) of the smallest model ($m = 1$). **(C)** Normalized residual heldout log-likelihood of models of size $n = 16$, as we increase the number of input channels. Here, we used 3 channels with the standard stimulus-dependent prior, and we used a flat prior for each additional channel. As in (A-B), normalization was done by subtracting the average residual log-likelihood (across animals, cells, and cross validation folds) of the smallest model ($m_{extra} = 0$). **(D)** Difference in the residual heldout log-likelhood of our standard models, and deterministic models driven by condition-dependent inputs that we set to be equal to the envelope of our prior. While we trained the deterministic models using the same objective as our standard models, their log-likelihood is considerably worse, as they are unable to capture any trial-to-trial variability. **(E)** Residual heldout log-likelihood of models trained with (left) and without (right) a bias term. For each setting, we selected the best model per animal. **(F)** The PV classification accuracy plotted against residual likelihoods for all trained models across all animals (each point is one model, 239 models overall). Error bars are $2 \times s.e.m.$ The marginal distributions are shown on top and right.

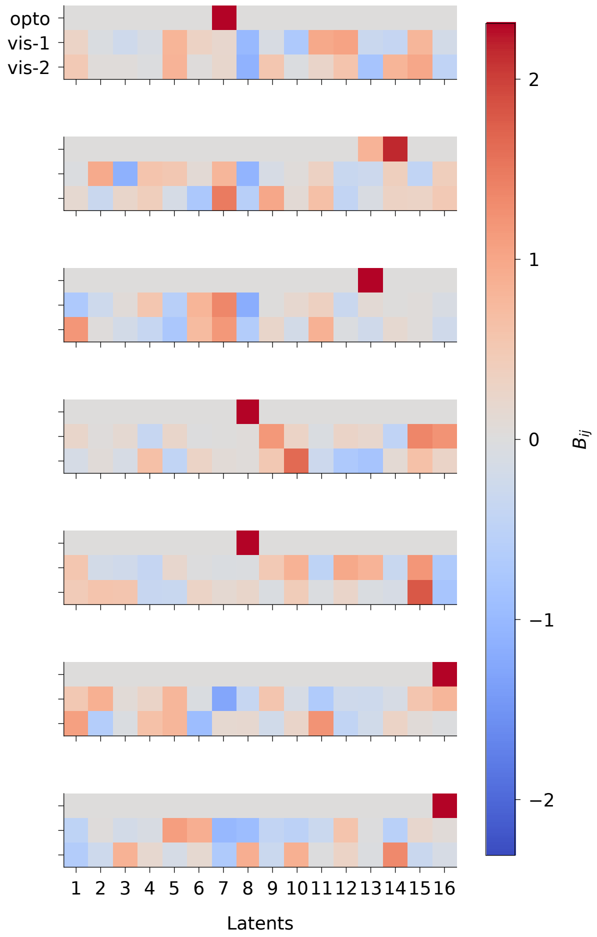


**Figure S2:** Visualization of the learned input coefficients ($\boldsymbol{B}$ matrices) for all animals. The rows correspond to the different input channels (opto and visual stimuli) and the columns to the $n = 16$ latent units.

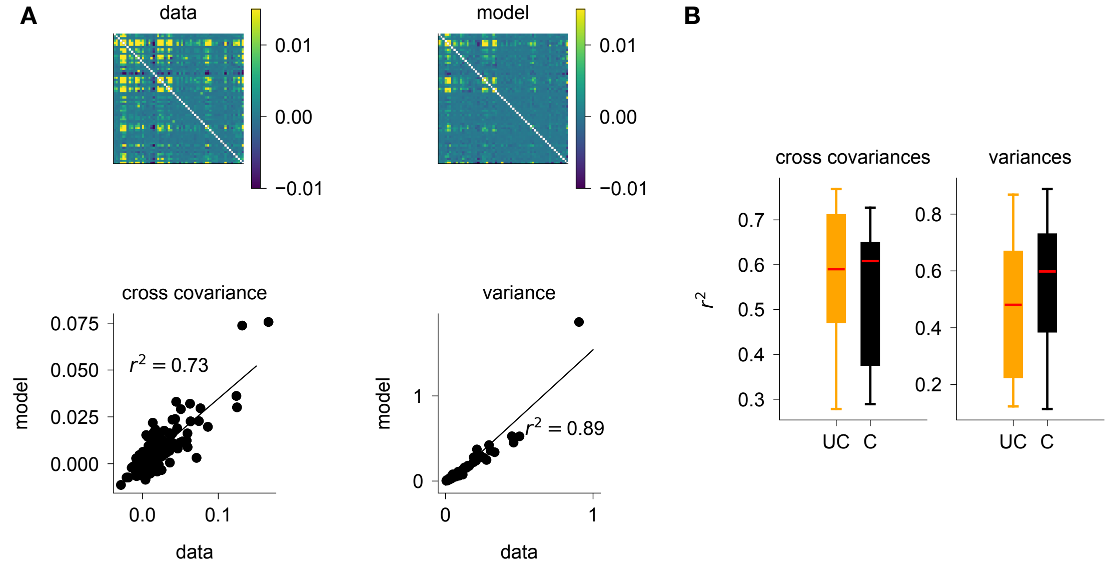


**Figure S3: Model performance on predicting covariances. (A)** Top: Covariance matrices of empirical neural firing (left) and model-predicted neural activity (right). Diagonal elements (variances) are not shown. Bottom left: Off-diagonal elements of the model predicted covariance matrix, plotted against the corresponding elements in the data covariance matrix, for one example animal. Bottom right: Empirical variance of neural activity plotted against the model-predicted ones, for the same example animal as in left. The model in (A) refer to the circuit-constrained model. **(B)** Distribution over animals of the coefficient of determination ($r^2$, model vs. data) for the off-diagonal elements of the covariance matrix (left) and the variances (right). The constrained models are shown in black and the unconstrained ones in orange.

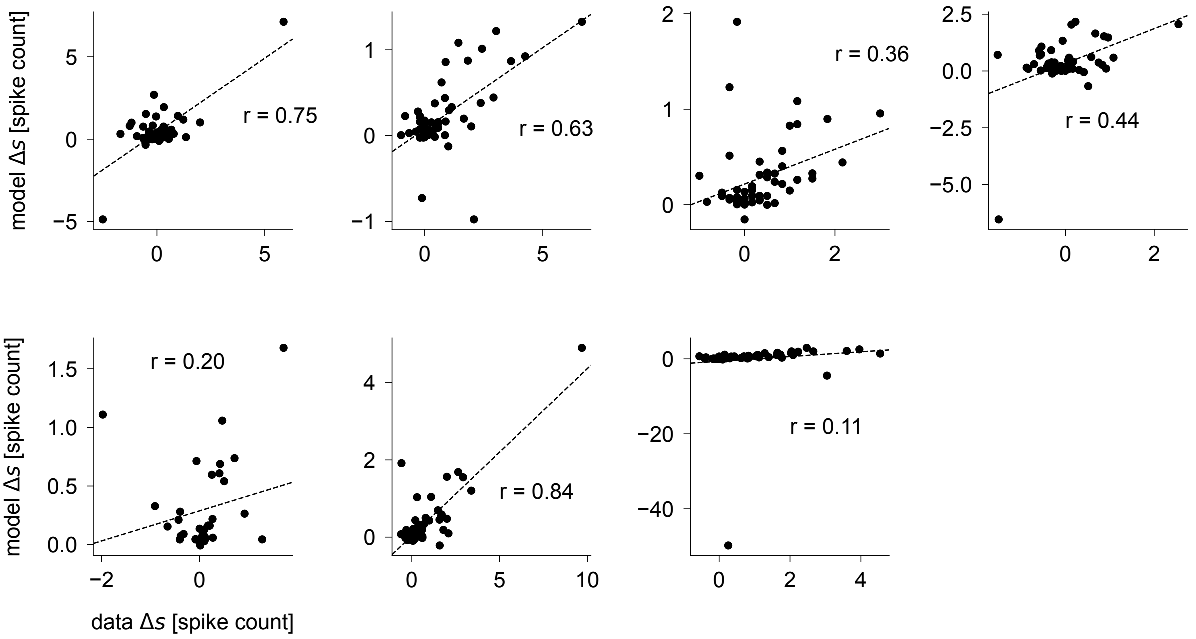


**Figure S4:** As in Figure 2D middle, but separately for each animal, using the same condition and optogenetic perturbation time window as the main figure.

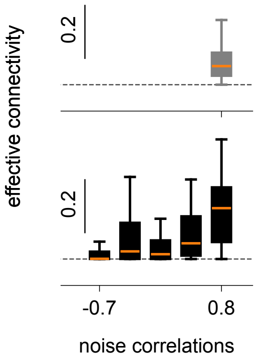


**Figure S5:** As in Figure 2G (main text), but using go instead of no-go trials

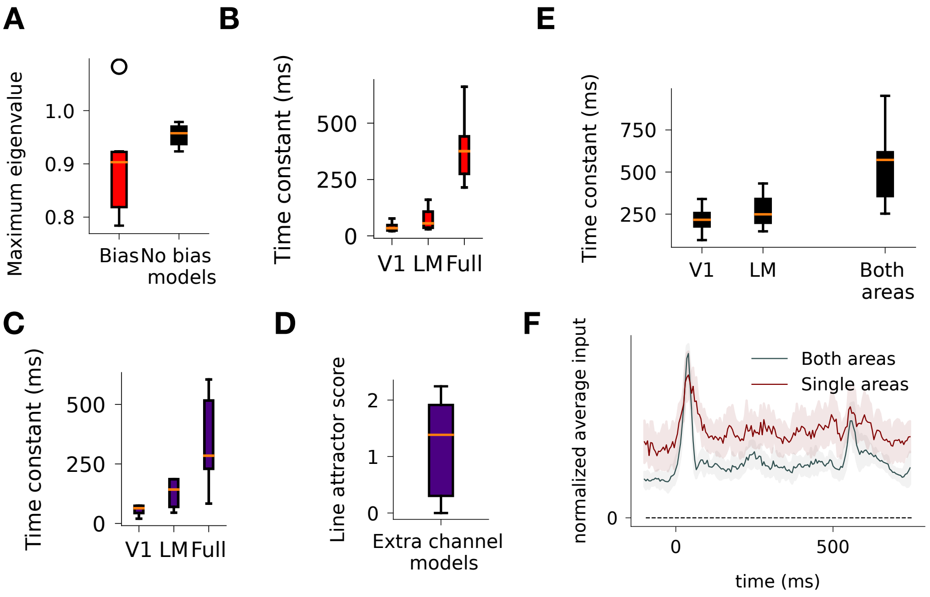


**Figure S6: Attractor controls (A)**: Comparison of the maximum eigenvalue of models whose bias was learned from the start (red), relative to models whose bias term was fixed to be **0** (black). We trained multiple models per animal (15 bias models and 50 no-bias models) and selected the one with the highest residual log-likelihood (see details in Methods). **(B)**: Distribution of the slowest timescale of individual areas, or the full dynamics, in models whose bias term was then allowed to be learned after 12000 iterations. For each animal, this analysis was performed for the model with the highest residual log-likelihood in the absence of bias (i.e the models analysed in the main text). **(C)**: Distribution of the slowest timescale of individual areas, or the full dynamics, in models with 1,2, or 3 additional input channels. For each animal, we trained 3 models per number of additional input channels, and selected the models that had the highest residual log-likelihood across that channel number and identified PV cells as inhibitory. This led to a total of $n = 9$ models across all animals. **(D)**: Distribution of line attractor scores of models with 1,2, or 3 additional input channels (selected in the same way as in C). **(E)**: Slowest timescale of the linearized dynamics in models fit to data from individual areas (V1 or LM), or to the whole network (Both areas). For each animal we fit models to the area that received optogenetic perturbation, and selected one model using the selection procedure described in Methods, yielding one model per animal. **(F)**: Average norm of the input to the latent circuit, for models fit to individual areas (selected in the same way as in E) in maroon, and models fit to the full circuit in gray. We normalize the norm to its maximal value across time for every trial, and take the average across trials for every animal. Line and shaded error bars denote the mean and standard error to the mean across models.

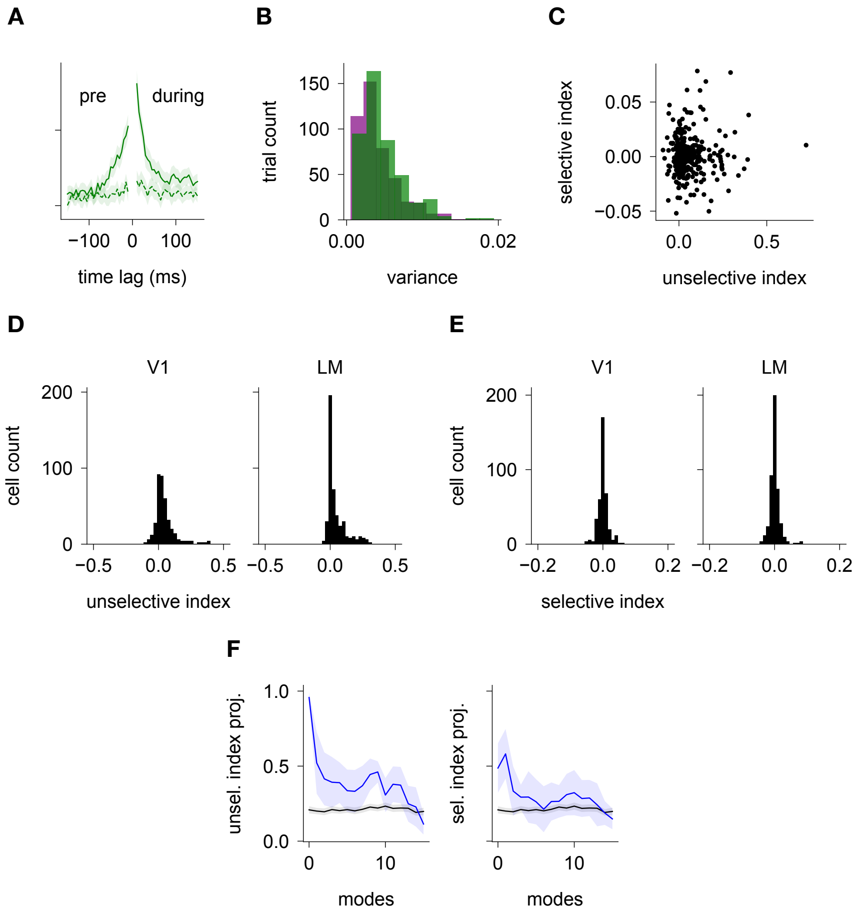


**Figure S7: (A)** Autocorrelation function of model predicted neural activity pre- (left half) and during stimulus (right half), projected onto the 'agree-balanced' (solid) and 'agree-unbalanced' (dashed) modes. **(B)** Distribution over trials of the total variance of neural activity during the stimulus, projected onto the 'agree-balanced' (green) and 'disagree-balanced' (purple) modes. Median difference (agree-disagree) = 0.0005, $p < 10^{-5}$ **(C)** Distribution of the selective indices across V1 and LM in all animals, against the nonselective indices. Pearson r = 0.05, p>0.05. **(D)** Distribution of the unselective indices across all neurons and animals, separately for V1 (left) ad LM (right) neurons (V1 mean = 0.046, $p < 10^{-5}$, LM mean = 0.047, $p < 10^{-5}$). P-values are two-sided t-tests. **(E)** Same as (D) but for selective indices (V1 mean = $2 \times 10^{-5}$, $p > 0.05$, LM mean = 0.0008, $p > 0.05$). **(F)** Alignment of the latent circuits' eigenmodes onto agree unselective (left) and selective modes (right) in blue (see Methods). The eigenmodes on the x-axis are ordered from the slowest (longest decay time constant) to fastest, and the y-axis shows the absolute value of the dot products between the eigenmodes and the unit-length vectors containing latent circuit's unselective or selective indices. The black lines denote chance level ($\pm$ C.I.), resulting from projecting random unit-length (16D) vectors onto the eigenmodes. The elements of the random vectors were drawn from a Gaussian distribution ($\mu = 0, \sigma = 1$), and 20 random vectors per animal were used to calculate confidence intervals.

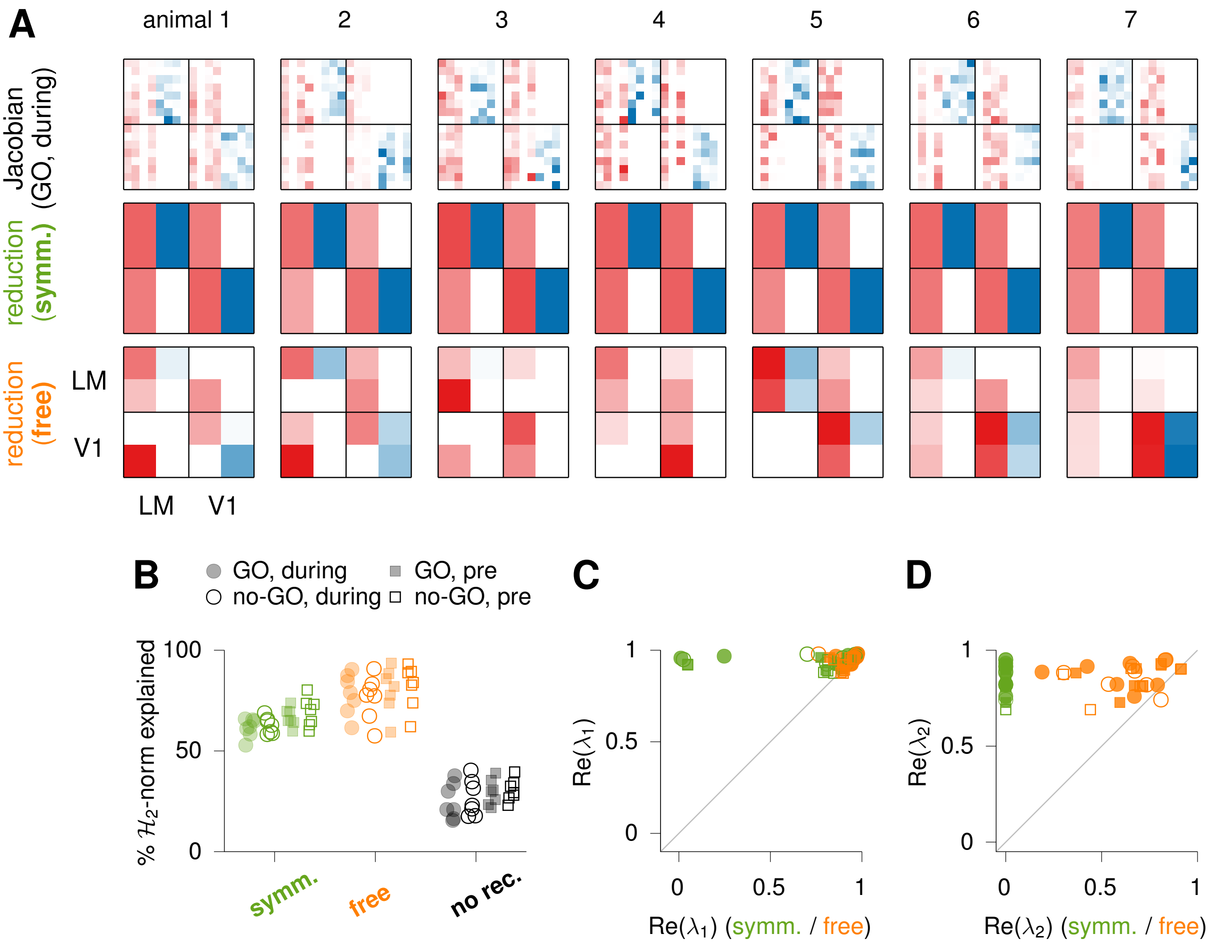


**Figure S8: Fitting minimal (4D) models to the learned dynamics**. (**A**) Full Jacobian (top; model linearized during presentation of the go stimulus) and best-fitting reduced linear model with (middle) and without (bottom) the structural symmetry constraints of Equation 33, for each animal (see Section S3). (**B**) Goodness of fit, calculated as the percentage of $\mathcal{H}_2$-norm of the full transfer function explained by the reduced model, shown here for all animals and all four Jacobian types (go/nogo stimulus, pre-and during stimulus presentation). Black symbols denote minimal models without recurrent connections, i.e. models in which only the input and output matrices are optimized. (**C**) Largest real part in the eigenvalue spectrum of the full Jacobians, against the largest real part in the spectrum of the reduced models. (**D**) Same as (C), for the second largest real part.

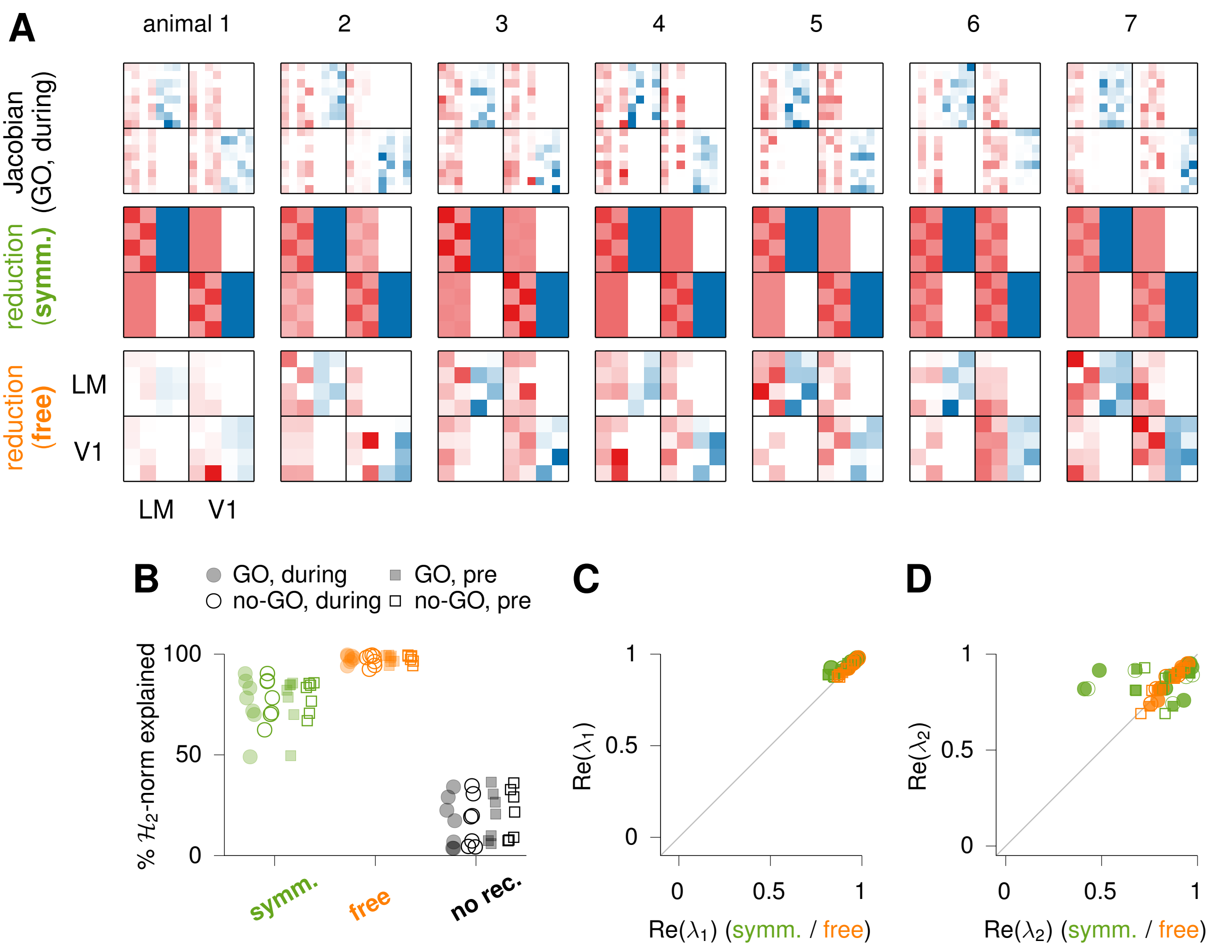

**Figure S9: Fitting minimal (8D) models to the learned dynamics**. Same as Figure S8 – caption repeated below for convenience. (**A**) Full Jacobian (top; model linearized during presentation of the go stimulus) and best-fitting reduced linear model with (middle) and without (bottom) the structural symmetry constraints of Equation 36, for each animal (see text). (**B**) Goodness of fit, calculated as the percentage of $\mathcal{H}_2$-norm of the full transfer function explained by the reduced model, shown here for all animals and all four Jacobian types (go/nogo stimulus, pre-and during stimulus presentation). Black symbols denote minimal models without recurrent connections, i.e. models in which only the input and output matrices are optimized. (**C**) Largest real part in the eigenvalue spectrum of the full Jacobians, against the largest real part in the spectrum of the reduced models. (**D**) Same as (C), for the second largest real part.

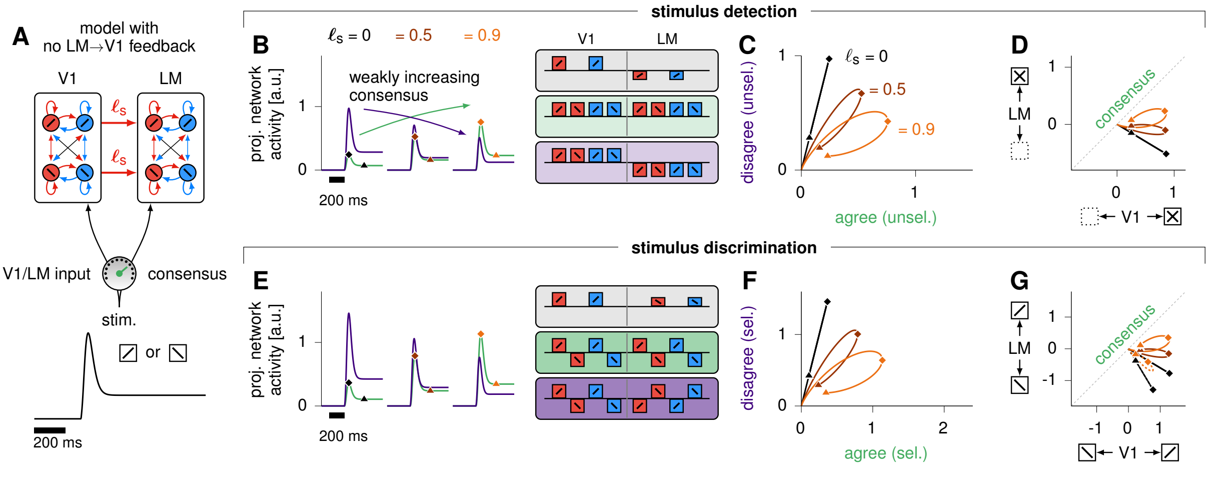

**Figure S10:** As in Figure 5 (main text), but with no feedback connections from LM to V1

### Supplementary notes

#### S1 Attractor controls

As mentioned in Discussion, one caveat with our approach is that it does not come with any identifiability guarantees. This means that there can be many models which fit the data equally well but display different dynamics or inputs. Of particular concern is the degeneracy that exists between dynamics and inputs, and the fact that, since we use a Gaussian prior over inputs, the model is implicitly encouraged to use as little inputs as possible to explain the data. We can therefore not rule out completely that the model may be hallucinating attractor dynamics, in order to explain the sustained activity while minimizing the “input cost”.

To assess the robustness of our findings, we therefore conducted several control experiments. Firstly, we evaluated the effect of incorporating a bias term in the dynamics, which can be used to explain the sustained activity without requiring any slow timescales in the dynamics. We found that such models were harder to train, as they were prone to falling into local minima early during training, which translated into a lower residual log-likelihood (Supplementary Material S1E). However, similarly to the main models, we observed that these models consistently relied on slow dynamics, with the eigenvalues of the linearized dynamics being close to, or higher than 1 (corresponding to slow or unstable dynamics; Supplementary Material S6A). Additionally, as the inclusion of the bias in the dynamics made models prone to falling into local minima, we evaluated the effect of pre-training the models without a bias term, and adding the bias to the trainable parameters after 12000 iterations. We found that this did not qualitatively change our results, with the models fine-tuned with bias still displaying dynamics much slower than the single time constant in the full network, and individual areas showing a much faster decay (see Figure S6B).

Another potential concern is that the attractor dynamics may emerge because of our main model’s limited number of input channels; to address this, we did the following controls. As shown in Figure S1, models with 3 channels had a higher log-likelihood than models with 4 or 5 channels (this held whether the additional channels had the stimulus prior or a flat prior). Additionally, attractor dynamics were also present in models with either 1,2, or 3 additional input channels (Figure S6C; see Figure S6D for the attractor score across models). Thus, while these models have more flexibility to use inputs to explain the sustained activity during stimulus presentation, we found that they nonetheless learned slow dynamics in the full network, and faster dynamics in individual areas.

Finally, to ensure that the attractor dynamics emerge from connecting the V1 and LM areas, and do not simply reflect a slow timescale in one of the two areas that the model attributes to long-range connections, we fit models to recordings from individual areas. For each animal, this corresponded to fitting two separate models to the V1 and LM recordings: importantly, the neurons in each of the two areas still exhibit sustained activity during the stimulus presentation. Interestingly, however, we found that those single-area models learned much faster dynamics than the full models (see Figure S6E). This was reflective of the fact that some of those models used more inputs during the stimulus period (see Figure S6F). This suggests that the slow dynamics do indeed emerge in the full network and not in individual areas.

Altogether, these controls suggest that the line attractor dynamics that we observe in our models are robust to specific modelling choices.

### S2 Minimal symmetric interconnected E/I networks with long-range excitatory connectivity

#### S2.1 Four-dimensional network

We considered E/I networks consisting of one E and one I neuron, described as a linear rate model, whose activity $\boldsymbol{r} = [r^{\mathrm{E}}, r^{\mathrm{I}}]^T$ in the absence of external input would evolves as:

$$\tau \dot{\boldsymbol{r}} = -\boldsymbol{r} + W\boldsymbol{r} \tag{S1}$$

$$W = \begin{bmatrix} e & -i \\ e & -i \end{bmatrix} \tag{S2}$$

Connecting two of these networks with excitatory long-range connections will result in the network in Figure 4 whose activity is described as:

$$\tau \dot{\boldsymbol{r}} = -\boldsymbol{r} + W\boldsymbol{r} = (W - I)\boldsymbol{r} = A\boldsymbol{r} \tag{S3}$$

where

$$\boldsymbol{r} = \begin{bmatrix} r_1^{\mathrm{E}} \\ r_1^{\mathrm{I}} \\ r_2^{\mathrm{E}} \\ r_2^{\mathrm{I}} \end{bmatrix} \quad \text{and} \quad W = \begin{bmatrix} e & -i & \ell & 0 \\ e & -i & \ell & 0 \\ \ell & 0 & e & -i \\ \ell & 0 & e & -i \end{bmatrix} \tag{S4}$$

where $\ell$ is the strength of long-range excitatory connectivity, and $I$ is the identity matrix. Using a similar approach to Murphy and Miller [2009], we define the following four patterns of activity that describe either E-I balanced or unbalanced activity which could either be in the same direction or opposite directions across the two areas.

$$\boldsymbol{b_a} = \frac{1}{2}\begin{bmatrix} 1 \\ 1 \\ 1 \\ 1 \end{bmatrix} \qquad \boldsymbol{u_a} = \frac{1}{2}\begin{bmatrix} 1 \\ -1 \\ 1 \\ -1 \end{bmatrix} \qquad \boldsymbol{b_d} = \frac{1}{2}\begin{bmatrix} 1 \\ 1 \\ -1 \\ -1 \end{bmatrix} \qquad \boldsymbol{u_d} = \frac{1}{2}\begin{bmatrix} 1 \\ -1 \\ -1 \\ 1 \end{bmatrix} \tag{S5}$$

Where $\boldsymbol{b}_a$ and $\boldsymbol{u}_a$ are the agreeing ('$a$') E-I balanced ('$b$') and unbalanced ('$u$') modes, and $\boldsymbol{b}_d$ and $\boldsymbol{u}_d$ are the disagreeing ('$d$') E-I balanced ('$b$') and unbalanced ('$u$') modes. The network activity projected on these vectors evolves as:

$$\tau \dot{\boldsymbol{b_a}} = A\boldsymbol{b_a} = (e - i + \ell - 1)\boldsymbol{b_a} \tag{S6}$$

$$\tau \dot{\boldsymbol{u_a}} = A\boldsymbol{u_a} = (e + i + \ell)\boldsymbol{b_a} - \boldsymbol{u_a} \tag{S7}$$

$$\tau \dot{\boldsymbol{b_d}} = A\boldsymbol{b_d} = (e - i - \ell - 1)\boldsymbol{b_d} \tag{S8}$$

$$\tau \dot{\boldsymbol{u_d}} = A\boldsymbol{u_d} = (e + i - \ell)\boldsymbol{b_d} - \boldsymbol{u_d} \tag{S9}$$

Furthermore these four vectors form an orthonormal basis, ($Q = [\boldsymbol{b_a}, \boldsymbol{u_a}, \boldsymbol{b_d}, \boldsymbol{u_d}]$) which is a Schur basis of A such that $\tilde{A} = Q^T A Q$ is an upper triangular matrix:

$$\tilde{A} = \begin{bmatrix} e - i + \ell - 1 & e + i + \ell & 0 & 0 \\ 0 & -1 & 0 & 0 \\ 0 & 0 & e - i - \ell - 1 & e + i - \ell \\ 0 & 0 & 0 & -1 \end{bmatrix} \tag{S10}$$

This upper triangular form describes feedforward connectivity in the new basis $Q$. Specifically, as seen in Figure S11, the network consists of two separate agreeing and disagreeing subnetworks, and each subnetwork corresponds to feedforward amplification from the unbalanced to balanced modes.

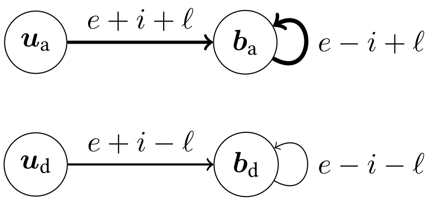


**Figure S11:** Equivalent feed-forward structure for two E/I networks connected via long-range excitatory connections.

The time constants of this network are determined by the eigenvalues of $\tilde{A}$ (which are also the eigenvalues of $A$), given by its diagonal values, i.e:

$$\lambda_{b_a} = e - i + \ell - 1 \tag{S11}$$

$$\lambda_{u_a} = -1 \tag{S12}$$

$$\lambda_{b_d} = e - i - \ell - 1 \tag{S13}$$

$$\lambda_{u_d} = -1. \tag{S14}$$

The unbalanced modes decay at the rate of the leak term, i.e with time constants $\frac{-\tau}{\lambda_{u_d}} = \frac{-\tau}{\lambda_{u_a}} = \tau$, where $\tau$ is the single neuron time constant.

In the balanced modes, decay time constants depend on $e$, $i$, and $\ell$. The time constant of the disagree-balance mode is $\tau_{d_b} = \frac{-\tau}{e-i-\ell-1}$. Since inter-area connections are excitatory, i.e. $\ell > 0$, $\frac{-\tau}{e-i-\ell-1} \leq \frac{-\tau}{e-i-1}$, meaning that activity in the disagree-balance mode decays faster than the time constant of the individual areas.

The agree-balanced mode, in contrast, has a time constant of $\frac{-\tau}{e-i+\ell-1} \geq \frac{-\tau}{e-i-1}$, corresponding to a decay slower than each individual area. In the approximately balanced network ($e \approx i$), this time constant grows without bound as $\ell \to 1$, corresponding to the emergence of an approximate line attractor along the agreeing-balanced mode.

#### S2.2 Orientation selective networks

##### S2.2.1 Local orientation selective network

We next considered an E-I network which incorporated selectivity to go or nogo stimuli ($45°$ or $-45°$ oriented stimulus). For this, we generalized the local network of Equation S1, such that instead of a single excitatory and a single inhibitory unit, we had excitatory and inhibitory *subpopulations*. We assumed two subpopulations of excitatory neurons and two subpopulations of inhibitory neurons, where one received direct go input, and the other receiving direct no-go input. We retained the assumption that the $W_{E\to I}$ and $W_{E\to E}$ connectivity matrices were the same and equal to $W_E$. Similarly, we defined $W_{I\to I} = W_{I\to E} = W_I$.

The connectivity within E and I subpopulations ($W_E, W_I$) had both stimulus selective ($e_s, i_s$), and unselective ($e, i$) components, such that :

$$W_E = \begin{bmatrix} e + e_s & e \\ e & e + e_s \end{bmatrix} \qquad W_I = \begin{bmatrix} i + i_s & i \\ i & i + i_s \end{bmatrix}. \tag{S15}$$

The local connectivity matrix in this network could therefore be written as:

$$W = \begin{bmatrix} W_E & -W_I \\ W_E & -W_I \end{bmatrix} \quad \text{and} \quad \boldsymbol{r} = \begin{bmatrix} r^{\mathrm{E}}_{\mathrm{go}} \\ r^{\mathrm{I}}_{\mathrm{go}} \\ r^{\mathrm{E}}_{\mathrm{nogo}} \\ r^{\mathrm{I}}_{\mathrm{nogo}} \end{bmatrix} \tag{S16}$$

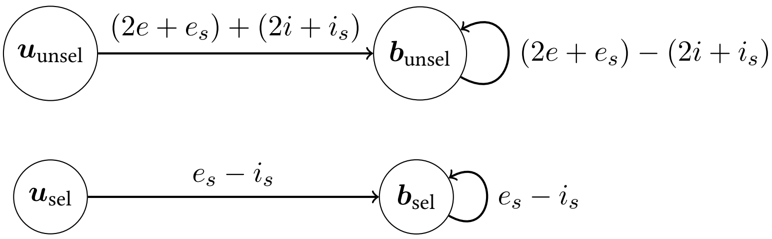


**Figure S12:** Feedforward equivalent of an E-I network with orientation selectivity.

Similarly to Section S2.1, it can be shown that $Q = \frac{1}{2}\begin{bmatrix} 1 & 1 & 1 & 1 \\ 1 & -1 & 1 & -1 \\ 1 & 1 & -1 & -1 \\ 1 & -1 & -1 & 1 \end{bmatrix} = \begin{bmatrix} \boldsymbol{b}^{\text{unsel}}, \boldsymbol{b}^{\text{sel}}, \boldsymbol{u}^{\text{unsel}}, \boldsymbol{u}^{\text{sel}} \end{bmatrix}$ is a Schur basis for $W$ and in this basis $W$ can be written as:

$$\tilde{W} = \begin{bmatrix} (2e + e_s) - (2i + i_s) & 0 & (2e + e_s) + (2i + i_s) & 0 \\ 0 & e_s - i_s & 0 & e_s + i_s \\ 0 & 0 & 0 & 0 \\ 0 & 0 & 0 & 0 \end{bmatrix} \tag{S17}$$

Again, in the Schur basis, the network can be expressed as two separate feedforward networks, each from an E-I unbalanced mode to an E-I balanced mode (Figure S12).

The feedforward weights are $(2e+e_s)+(2i+i_s)$ for the unselective network and $e_s+i_s$ for the selective network. Similar to Section S2.1, the decay time constants can be calculated from the eigenvalues of $\tilde{A} = \tilde{W} - I$. Both balanced modes decay with a time constant of $\tau$ and the unselective and selective balanced modes have a time constant of $\tau_{unsel} = \frac{-\tau}{2(e-i)+(e_s-i_s)-1}$ and $\tau_{sel} = \frac{-\tau}{(e_s-i_s)-1}$.

##### S2.2.2 Two connected orientation selective networks

Next, we moved to a configuration where two selective networks, as described in section S2.2.1, were connected with long-range excitatory connectivity matrix $W_L$. The connectivity of this network was described by:

$$W = \begin{bmatrix} W_E & -W_I & W_L & 0 \\ W_E & -W_I & W_L & 0 \\ W_L & 0 & W_E & -W_I \\ W_L & 0 & W_E & -W_I \end{bmatrix} \quad \text{and} \quad \boldsymbol{r} = \begin{bmatrix} r_1^{\text{E,go}} \\ r_1^{\text{E,nogo}} \\ r_1^{\text{I,go}} \\ r_1^{\text{I,nogo}} \\ r_2^{\text{E,go}} \\ r_2^{\text{E,nogo}} \\ r_2^{\text{I,go}} \\ r_2^{\text{I,nogo}} \end{bmatrix} \tag{S18}$$

where $W_L$ is also a $2 \times 2$ matrix, structured similarly to $W_E$ and $W_I$:

$$W_L = \begin{bmatrix} \ell + \ell_s & \ell \\ \ell & \ell + \ell_s \end{bmatrix}. \tag{S19}$$

Therefore, the full connectivity matrix can be written as:

$$W_{\text{full}} = \begin{bmatrix} e+e_s & e & -(i+i_s) & -i & \ell+\ell_s & \ell & 0 & 0 \\ e & e+e_s & -i & -(i+i_s) & \ell & \ell+\ell_s & 0 & 0 \\ e+e_s & e & -(i+i_s) & -i & \ell+\ell_s & \ell & 0 & 0 \\ e & e+e_s & -i & -(i+i_s) & \ell & \ell+\ell_s & 0 & 0 \\ \ell+\ell_s & \ell & 0 & 0 & e+e_s & e & -(i+i_s) & -i \\ \ell & \ell+\ell_s & 0 & 0 & e & e+e_s & -i & -(i+i_s) \\ \ell+\ell_s & \ell & 0 & 0 & e+e_s & e & -(i+i_s) & -i \\ \ell & \ell+\ell_s & 0 & 0 & e & e+e_s & -i & -(i+i_s) \end{bmatrix} \tag{S20}$$

Combining insights from Section S2.1 and Section S2.2.1, we found the Schur basis of $W$ ($Q_{\text{full}}$), by stacking the Schur basis of the single-area selective network $Q$ in the following arrangement:

$$Q_{\text{full}} = \frac{1}{\sqrt{2}} \begin{bmatrix} Q & Q \\ Q & -Q \end{bmatrix} = [\boldsymbol{b}_a^{\text{unsel}}, \boldsymbol{b}_a^{\text{sel}}, \boldsymbol{u}_a^{\text{unsel}}, \boldsymbol{u}_a^{\text{sel}}, \boldsymbol{b}_d^{\text{unsel}}, \boldsymbol{b}_d^{\text{sel}}, \boldsymbol{u}_d^{\text{unsel}}, \boldsymbol{u}_d^{\text{sel}}] \tag{S21}$$

Where ‘b’/‘u’ refer to ‘balanced’/‘unbalanced’ modes, ‘a’/‘d’ refer to ‘agree’/‘disagree’ modes, and ‘unsel’/‘sel’ refer to ‘unselective’/‘selective’ modes.

In the Schur basis $Q_{\text{full}}$, the full connectivity matrix $W_{\text{full}}$ is expressed as:

$$\begin{bmatrix} w^- + (2\ell+\ell_s) & 0 & w^+ + (2\ell+\ell_s) & 0 & 0 & 0 & 0 & 0 \\ 0 & w_s^- + \ell_s & 0 & w_s^+ + \ell_s & 0 & 0 & 0 & 0 \\ 0 & 0 & 0 & 0 & 0 & 0 & 0 & 0 \\ 0 & 0 & 0 & 0 & 0 & 0 & 0 & 0 \\ 0 & 0 & 0 & 0 & w^- - (2\ell+\ell_s) & 0 & w^+ - (2\ell+\ell_s) & 0 \\ 0 & 0 & 0 & 0 & 0 & w_s^- - \ell_s & 0 & w_s^+ - \ell_s \\ 0 & 0 & 0 & 0 & 0 & 0 & 0 & 0 \\ 0 & 0 & 0 & 0 & 0 & 0 & 0 & 0 \end{bmatrix}$$

where

$$w^- = (2e+e_s) - (2i+i_s) \tag{S22}$$
$$w^+ = (2e+e_s) + (2i+i_s) \tag{S23}$$
$$w_s^- = e_s - i_s \tag{S24}$$
$$w_s^+ = e_s + i_s \tag{S25}$$

Therefore, similarly to the non-selective network of Section S2.1, the network decomposes into an agree and a disagree network. However, here, each network has a selective and an unselective subnetwork. The time constants of all four unbalanced modes are $\tau$ as before, and the balanced modes evolve with the following constants: In the agree network:

$$\tau_{b_a}^{\text{unsel}} = \frac{-\tau}{2(e-i) + (e_s - i_s) + (2\ell + \ell_s) - 1} \tag{S26}$$
$$\tau_{b_a}^{\text{sel}} = \frac{-\tau}{(e_s - i_s) + \ell_s - 1} \tag{S27}$$

And in the disagree network:

$$\tau_{b_d}^{\text{unsel}} = \frac{-\tau}{2(e-i) + (e_s - i_s) - (2\ell + \ell_s) - 1} \tag{S28}$$
$$\tau_{b_d}^{\text{sel}} = \frac{-\tau}{(e_s - i_s) - \ell_s - 1} \tag{S29}$$

Given that $\ell, \ell_s \geq 0$, connecting the two areas results in slowing down both selective and unselective agreeing networks and speeding up both disagreeing networks. If inter-area connectivity is completely non-selective or untuned, that is $W_L = \begin{bmatrix} \ell & \ell \\ \ell & \ell \end{bmatrix}$, then $\ell_s = 0$ and inter-area connectivity only affects the unselective activity modes (slows down unselective-agree and speeds up unselective-disagree modes). However, if inter-area connectivity is tuned, then both the unslective and selective networks get affected. And the degree to which the agree-selective network slows down depends on the specificity of long-range connections (if there is no untuned long-range connectivity, $\ell = 0$, then the unselective and selective networks slow down to the same degree).

#### S3 Learned dynamics are well approximated by the minimal models

In Figures S8 and S9, we assessed the degree to which the minimal models of Figure 4 could quantitatively capture the dynamics of the latent circuit models we had obtained from data. We did this for both the unselective (Figure 4B) and selective (Figure 4C) minimal models. For each minimal model class, we optimized the parameters of the minimal model ('reduced model') to capture as much of the impulse response of the linearized latent circuit ('full model'; see details below).

**Fitting 4D (unselective) minimal models** Each full model was characterized by (i) a matrix of recurrent weights $W \in \mathbb{R}^{16\times16}$ and (ii) a matrix of input weights $B \in \mathbb{R}^{16\times3}$. The best we could expect from the 4-dimensional reduced models was that they matched the *mean* impulse responses in local E and I populations in the full model. Therefore, we constructed a readout (output) matrix

$$C = \frac{1}{4}\begin{bmatrix} 1 & \dots & 1 & & & & & & & & & \\ & & & 1 & \dots & 1 & & & & & & \\ & & & & & & 1 & \dots & 1 & & & \\ & & & & & & & & & 1 & \dots & 1 \end{bmatrix} \in \mathbb{R}^{4\times16} \tag{S30}$$

where all unspecified elements are zeros. Accordingly, each reduced model was characterized by (i) a matrix of recurrent weights $W_{\text{red}} \in \mathbb{R}^{4\times4}$, (ii) a matrix of input weights $B_{\text{red}} \in \mathbb{R}^{4\times3}$, and (iii) a matrix of output weights $C_{\text{red}} \in \mathbb{R^+}^{4\times4}$. The discrepancy between the impulse responses of the full and reduced input-output systems was then computed as follows (see also Kao and Hennequin, 2019). Consider the residual system $(\hat{W}, \hat{B}, \hat{C})$ with

$$\hat{W} \equiv \begin{bmatrix} W & 0 \\ 0 & W_{\text{red}} \end{bmatrix} \qquad \hat{B} \equiv \begin{bmatrix} B \\ B_{\text{red}} \end{bmatrix} \qquad \hat{C} \equiv \begin{bmatrix} C & -C_{\text{red}} \end{bmatrix} \tag{S31}$$

If the reduced model fully captured the local mean responses of the full model, then the responses of this residual system to an impulse in any of the three input channels would be zero. Thus, a sensible way of quantifying the accuracy of the model reduction is to use the $\mathcal{H}_2$-norm of the residual system:

$$\mathcal{H}_2(\hat{W}, \hat{B}, \hat{C}) \equiv \text{Tr}(\hat{C}P\hat{C}^\top) \quad \text{with} \quad (\hat{W} - I)P + P(\hat{W} - I)^\top + \hat{B}\hat{B}^\top = 0 \tag{S32}$$

We used Adam to perform gradient-based minimization of Equation S32 w.r.t. the parameters of the reduced model, i.e. the parameters that determine $(W_{\text{red}}, B_{\text{red}}, C_{\text{red}})$ (see below). For this, we relied on automatic differentiation of Lyapunov solvers [Kao and Hennequin, 2020]. The quality of fit metric reported in the main text is defined as

$$1 - \frac{\min \mathcal{H}_2(\hat{W}, \hat{B}, \hat{C})}{\mathcal{H}_2(W, B, C)}.$$

The parameters of the reduced model were (i) the strength of local excitation ($e$ in Equation S4), (ii) the matrix of input weights $B_{\text{red}}$ with the third column (photo-stimulation input channel) constrained to be zero for all subpopulations except for the inhibitory subpopulation known to express ChR2 in the experiments, and (iii) the matrix of output weights $C_{\text{red}}$ which we constrained to be positive only. For the other parameters of the reduced recurrent weights, namely the strengths of local inhibition $i$ and long-range excitation $\ell$ (c.f. Equation S4), we directly set them to match $e/i$ and $\ell/e$ to the equivalent ratios in the full model. This led to more consistent model reductions without impacting the residual $\mathcal{H}_2$-norm.

For the 'free' model reductions (Figure S8, orange), we optimized all possible recurrent weights independently, whilst still respecting the E/I sign constraints as well as the absence of long-range I connections. We also compared with reduced models that have no recurrent connections at all (Figure S8, black, 'no-rec.').

**Fitting 8D (selective) minimal models** The approach we took for fitting the selective minimal models was analogous to the one used for the 4D model reductions (c.f. above), with a few differences:

- For these 8D models, we could expect the reduced model to capture not only the mean (unselective) responses in local E and I populations in the full model, but also the (selective) responses of smaller sub-populations therein, obtained by partitioning neurons locally according to their preference for either the $+45°$ or $-45°$ stimulus. We used the inferred latent responses of the full model in the different stimulus conditions (0-150 ms) to label each latent unit in the full model as preferring one or the other stimulus. We then introduced a matrix of positive output weights $C$ for the full model, which was structured similarly to Equation S30: it had 8 rows, each structured such that it decoded a weighted average responses of one of those selective sub-populations ($8 = 2$ (V1 vs. LM) $\times 2$ (E vs. I) $\times 2$ ($+45°$ vs. $-45°$)). We optimized all non-zero weights of this matrix $C$, subject to positivity constraints, along with the parameters of the reduced model, to minimize the $\mathcal{H}_2$-norm of the residual as detailed above.
- the $B_{\text{red}}$ matrix now had size $8 \times 3$
- the recurrent weights $W_{\text{red}}$ of the reduced model were as described in Equation S20, with the following re-parameterization:

$$i = e\alpha\sqrt{\frac{1+(1+\gamma_{\text{E}})^2}{2}} \qquad e_s = e\gamma_{\text{E}} \qquad \ell_s = e\beta\sqrt{\frac{1+(1+\gamma_{\text{E}})^2}{1+(1+\gamma_{\ell})^2}} \tag{S33}$$

  where $e$, $\gamma_{\text{E}}$ and $\gamma_{\text{I}}$ are positive free parameters, $\alpha$ is the overall ratio of local inhibition to local excitation in the full latent circuit, and $\beta$ is the overall ratio of long-range over local excitation in the full latent circuit. In other words, this parameterization enabled flexibility in the overall magnitude of the connectivity ($e$), as well as degree of selectivity in local and long-range excitatory connections ($\gamma_{\text{E}}$ and $\gamma_{\ell}$), but otherwise respected the above-mentioned ratios. Giving the model more flexibility by relaxing these ratio constraints did not improve the quality of fit, but resulted in more degenerate solutions.

Ta-Chu Kao and Guillaume Hennequin. Automatic differentiation of Sylvester, Lyapunov, and algebraic Riccati equations. *arXiv preprint arXiv:2011.11430*, 2020.
